# Chimpanzee culture beyond the conspicuous: Evidence for broad-scale observational social learning in wild individuals

**DOI:** 10.1101/2025.10.28.681847

**Authors:** Nora E. Slania, Mariana Gómez-Muñoz, Ayrin-Sophie Piephoh, Geresomu Muhumuza, Richard Young, T. Revathe, Catherine Hobaiter, Klaus Zuberbühler, Caroline Schuppli

**Affiliations:** Development and Evolution of Cognition Research Group, Max Planck Institute of Animal Behavior, Konstanz, Germany; Department of Comparative Cognition, Institute of Biology, University of Neuchâtel, Neuchâtel, Switzerland; Budongo Conservation Field Station, Masindi, Uganda; Wild Minds Lab, School of Psychology and Neuroscience, University of St Andrews, St Andrews, UK

**Keywords:** Animal Culture, Primates, Apes, Skill Acquisition, Social Transmission, Peering, Cultural Traits, Behavioral Repertoire, Feeding Skills, Diet repertoire

## Abstract

Wild chimpanzees possess diverse cultural repertoires, representing the richest example of non-human animal cultures. However, traditional methods, which investigate culture at the group level, have likely underestimated the full extent of chimpanzee cultural repertoires. In particular, the cultural relevance of everyday behaviors has remained largely unexplored. Here, we investigated evidence for social transmission of everyday behaviors to assess the breadth of individuals’ cultural repertoires in a population of wild eastern chimpanzees (*Pan troglodytes schweinfurthii*). First, we validated whether peering (i.e. close-range observation of a conspecific) serves as an indicator of social (i.e., cultural) learning. We then examined the contexts in which chimpanzees engage in peering to determine the range of behaviors that may be culturally transmitted. Finally, we explored potential motivations and additional functions of peering behavior. Our results indicate that chimpanzees use peering for targeted social information seeking in learning-intensive contexts. Peering rate was highest during immaturity, for complex or rare food items, and when observing older, more experienced conspecifics. Overall, wild chimpanzees peered at a wide range of everyday skills, such as feeding and grooming, and directed peering towards various conspecifics from an early age. We found no evidence supporting peering as a begging or submissive “gesture”, but our findings indicate that it may function as a signal to initiate affinitive interactions. Our findings suggest that wild chimpanzees use peering to learn a broad variety of skills, thereby highlighting unrecognized cultural potential in everyday skills. Furthermore, our findings suggest that peering may have multiple functions and underlying motivations.

## Introduction

Animal cultures are being recognized in an increasingly wide range of species^1^. While early research was motivated by the recognition of individual cultural traits^2–4^ — i.e. specific behaviors that are passed on within and between generations through social learning^5^ — studies on non-human primates, in particular apes, have since highlighted the potential for diverse and broad cultural repertoires^6–9^. The most striking and well-researched evidence for non-human primate culture are behavioral differences across wild populations, as evident in the presence and absence of behaviors^10–12^, as well as differences in the forms of behaviors^13–15^. Because these differences cannot be explained by ecological or genetic factors, they are most likely the result of individual innovation and subsequent social transmission of the innovated behavior^10^. Different forms of social learning may underlie cultural transmission, likely constituting a continuum, ranging from low-fidelity social learning via local enhancement or social facilitation to high-fidelity forms such as learning through direct observation of or interaction with a conspecific^16,17^. Social transmission and forms of social learning have been investigated across a narrow range of contexts^18–33^, such as the socially mediated spread of moss-sponging^33^ or acquisition of nut cracking^34–37^ in wild chimpanzees. However, estimates of the breadths of non-human primate cultures are commonly derived from group-specific behavioral repertoires^6–9,38,39^. These group-level comparisons – while offering striking evidence for the existence of non-human cultures^40^ – are prone to overlook socially transmitted traits where they do not result in conspicuous group-level differences^38,41^: Socially learned behaviors might share important characteristics under similar environmental conditions or might covary with ecological differences. Additionally, socially learned skills might be overlooked when they are inconspicuous, such as common feeding skills^41^, or show subtle variation within recognized techniques, such as chimpanzee termite fishing^38^. In this way, traditional approaches to non-human primate culture have likely underestimated the extent of socially acquired skills and hence of potential cultural traits within groups^41,42^.

Across non-human primates, chimpanzees (*Pan troglodytes*) provide the most detailed reports of cultural diversity^43,44^. Several decades of research have mapped and identified vast cultural repertoires across wild communities. Candidate cultural behaviors are commonly conspicuous^32,45,46^ and entail multi-step behavioral sequences (henceforth called complex behaviors or skills)^47–50^. Likewise, studies investigating wild chimpanzee skill acquisition have focused on these complex skills, many of which include the use of tools^18–31^. This body of research has found multiple lines of evidence that young chimpanzees learn complex skills through observing or interacting with role models (e.g. through begging and scrounging of food and foraging tools). Both observational and interactive forms of learning are assumed to be more efficient than more indirect types of social learning (such as forms of enhancement or facilitation) or individual learning, as they increase the signal-to-noise ratio of information and allow learners to gain access to more detailed information^51–53^. Learning through observation or interaction thus allows for more targeted exploration, minimizing risks of bodily harm associated with exploration, such as injuries or ingesting poisonous foods^51–53^.

In contrast to interactive forms of social learning, observational forms of social learning require lower levels of social tolerance and pose lower risks (e.g. from aggression or infectious disease transmission) because they do not require physical contact between the learner and conspecific^54,55^. Accordingly, direct observation of conspecifics has frequently been proposed as a form of learning in complex skill acquisition contexts (henceforth called observational social learning)^18,19,25,26,28,31,32^. Matsuzawa et al.^34^ termed learning via observation of conspecifics “education by master-apprenticeship” and suggested that young chimpanzees rely greatly on the close observation of others to develop skills. Systematic studies on orangutans^56–58^ (*Pongo abelii*) and capuchins^59^ (*Cebus capucinus*) have found multiple lines of evidence that individuals use the close-range attentive observation of a conspecific, i.e. peering^56^ (Figure 1), as a means to seek social information, which can result in learning. While observations of conspecifics are frequently reported in wild chimpanzees ^18,19,25,26,28,31,32^, and captive studies have outlined their potential to learn through observation^60–62^, systematic evidence validating peering as a behavioral indicator of targeted information seeking and subsequent learning in chimpanzees has only recently begun to emerge. So far, this evidence has narowly focused on complex skills, like tool use^28,63^.

**Figure 1:**
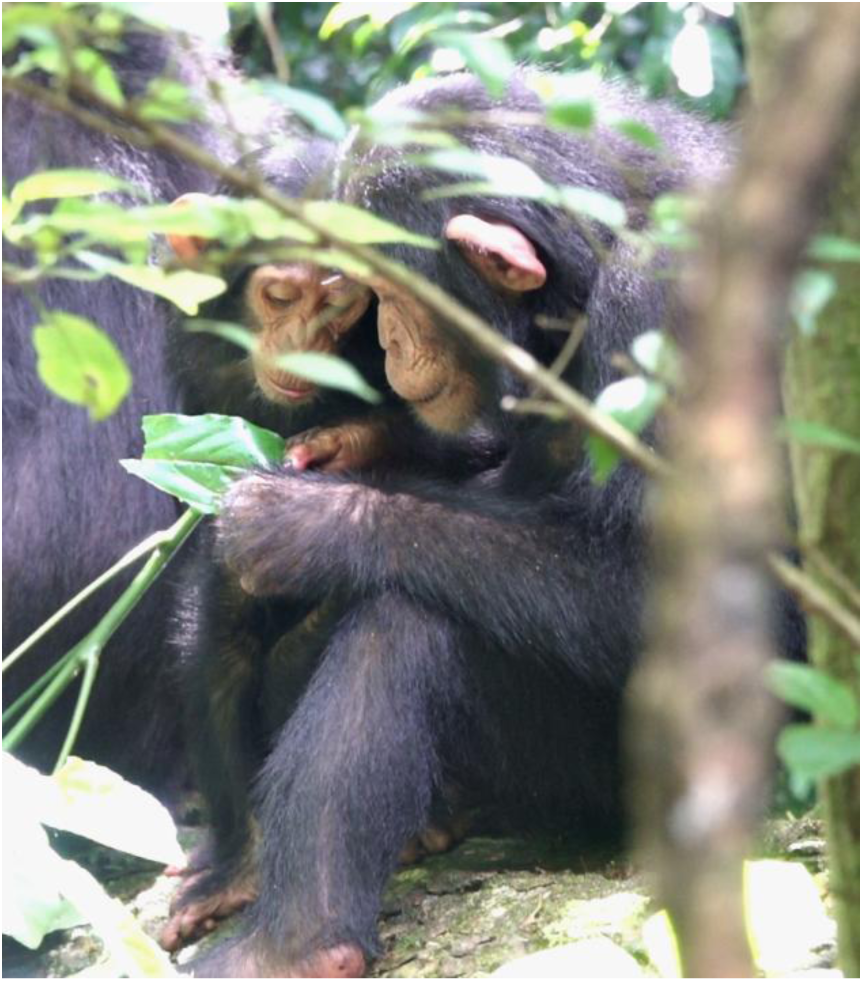
Infant chimpanzee (left) peering at the hands of a juvenile (right) engaging in ectoparasite inspection with a leaf. Picture: Nora E. Slania

The behaviors for which observational social learning and/or cultural variation have been studied tend to make up only a fraction of individuals’ activity budgets (e.g., in most chimpanzee populations, only a small fraction of the diet is acquired via tool use^31,48,64–66^). Everyday behavioral repertoires are composed of inconspicuous, simple skills, such as feeding techniques and social interactions comprising a small number of behavioral steps only. These routine repertoires are central to chimpanzee survival and encompass a broad range of knowledge and skills. For example, an adult chimpanzee’s feeding repertoire includes knowledge of the edibility and feeding techniques needed for often several hundred food items^67^. Given that immatures have to acquire behavioral repertoires within certain developmental timeframes^68^ and might draw fitness benefits from acquiring their repertoires quickly^69^, it is likely that observational social learning is widely used in everyday skill acquisition. Accordingly, in other primates, studies have found evidence for targeted social information seeking during skill acquisition of everyday skills such as feeding and nest building^56–59,70^. To date, wild chimpanzees’ everyday skill acquisition remains under-investigated, including the extent to which it is mediated through observational social learning. At the same time, the currently recognized repertoire of cultural behaviors in chimpanzees, although likely an underestimation, represents the largest known catalog of non-human animal culture. Hence, understanding how chimpanzees use observational social learning in daily life will advance our understanding of the true breadth of chimpanzee and other animal cultures. While learning can occur throughout life, studying immature individuals is particularly revealing, as they must acquire a vast variety of skills—many of which are simple, inconspicuous, and common, and may show limited individual variation in adulthood—making their skill acquisition a promising window into a species’ cultural scope.

As targets for observational learning, i.e. role models, ape mothers have received particular attention. Several studies on complex skill acquisition in chimpanzees highlight their importance as role models for young infants^18–23,26,30,31,71^. Whiten and de Waal^72^ proposed that social information seeking from the mother constitutes a first phase of social learning, which is later followed by a second phase of learning defined by increasing attention towards other group members. For chimpanzees, there is evidence that the multi-year acquisition of complex skills follows these phases (e.g., nut-cracking skill acquisition^71^). Notably, in this context, observational social learning regularly co-occurs with interactive forms of learning, which require more tolerant role models (see above). It has, however, remained unclear to what extent these patterns apply to the acquisition of simpler, i.e. less learning-intensive, everyday skills. As these contexts might require lower levels of social tolerance, and as immature chimpanzees are commonly surrounded by multiple conspecifics, chimpanzees might learn everyday skills from a wide range of individuals from a young age.

The first objective of this study is to investigate the validity of peering as a behavioral indicator of observational social learning in chimpanzees. We therefore investigate whether peering is used for targeted social information seeking and occurs in contexts in which learning is expected. We predict 1a) high peering rates during immaturity when individuals are acquiring their skill sets but very low rates during adulthood^56,59^; 1b) higher peering rates with increasing food item complexity and rarity^56,73^; 1c) chimpanzees to direct most peering at skilled individuals, i.e. older or similarly aged group members, but not at less skilled, i.e. younger individuals^58,59,71^.

The second objective of this study is to critically expand the common research focus, typically centred on complex and conspicuous skills, by investigating the extent to which chimpanzees use peering across their broader behavioral repertoire and to understand from whom individuals seek information. We predict 2a) peering target activities to comprise a wide range of behaviors, including skills beyond tool and object use; 2b) frequent peering at routine skills, such as grooming and feeding; 2c) peering across the whole ranges of food item complexity and rarity (see prediction 1b); 2d) diverse role model selection from a young age onwards.

While studies have convincingly linked peering to social information seeking (see above), it is plausible that this is neither its sole evolved function nor its immediate underlying motivation. Instead, peering could for example be motivated by a want to access an item and correspondingly function as a begging gesture (initially suggested in bonobos, *Pan paniscus*^74^). Peering could further serve to signal submissiveness to a more dominant individual, as has also been suggested in bonobos^75^. Lastly, particularly in social settings, peering could constitute a way to signal interest to engage in or join a social activity such as playing or grooming (henceforth called initiating affinitive interactions, following^76^). Importantly, these proposed functions, including peering as a means of social information seeking, are not mutually exclusive. Instead, diverse drivers behind peering behavior could create a broad scope of contexts in which direct observation allows for knowledge transfer, therefore promoting cultural developments across behavioral settings. Investigating diverse drivers of and motivations underlying peering may thus yield important novel insights into the dynamics of cultural processes.

Thus, the third objective of this study is to assess whether peering serves additional functions in everyday chimpanzee lives. Hence, we 3a) investigate whether peering is a form of begging behavior. We predict active begging behaviors (see below) to result in item transfer, but much less when peering alone. Further, if chimpanzees use peering to signal submission, we 3b) predict peering to be most frequently directed at high-ranking individuals and, further, in adult individuals to follow patterns of dominance in hierarchy. Lastly, to test whether peering is used to initiate affinitive interactions, we 3c) investigate how frequently peering is followed by affinitive interactions, in particular whether grooming partners show increased daily grooming rates when peering takes place.

We collected data on eastern chimpanzees (*Pan troglodytes schweinfuthii*) of the Sonso community in the Budongo Central Forest Reserve, Uganda. The Sonso chimpanzees are one of the few known chimpanzee communities that do not engage in any stick tool use and generally show relatively low levels of tool and object use^10,77^, allowing us to study the potential use of observational social learning where it has so far been overlooked. Our data collection comprised daily follows on instantaneous scan sampling at 3-minute intervals of activities of a focal individual and conspecifics within close proximity (<5m), as well as all occurrence focal data on peering and active begging. We analyzed ∼1,100h of focal follow data collected over ∼2.5 years on 28 focal individuals (17 immatures, 11 adults).

## Results

Using a Bayesian approach, we fitted five generalized linear mixed models (GLMMs) with negative binomial error structure to evaluate our predictions of how peering frequencies are affected by focal age, food item complexity (i.e. number of required pre-ingestive processing steps), and food item frequency (i.e. frequency of the item in the overall population diet) (Model A), by conspecific age (Model B), by kin relatedness (Model C1 and C2), and by conspecific rank (Model D). The number of peering events constituted the response variable for these models, and we included offset-terms to account for hours of opportunities (Model A, B, C1, D) or observation (Model C2). In addition, we fitted a GLMM with beta error structure (Model E) to assess whether grooming rates of individual dyads increased on days when one dyad partner had peered at the other dyad partner’s grooming behaviour.

### Model A: Effects of focal age, food item frequency, and food item complexity on peering (predictions 1a-b, 2c)

Peering rates changed with peerer age and were affected by the complexity and frequency of the food item processed by the peering target. We found that peering initially increased with age, peaking when the focal was around 5.4 years old for average food item complexity and frequency, and thereafter gradually decreased until early adulthood (age estimate=-3.25 [-2.39, -0.6], age^2^ estimate=-3.17 [-5.47, -1.19]), with no peering occurring later in life. Peering rates increased with decreasing food item frequency (estimate=-0.7 [-1.27, -0.09]) and with increasing food item complexity (estimate=0.64 [0.25, 1.08]), Figure 2. The overall model could account for 74% [58%, 83%] of the variation in peering rates. See Table S1 for model output and Figure S2 for Highest Density Intervals (HDI)s.

**Figure 2:**
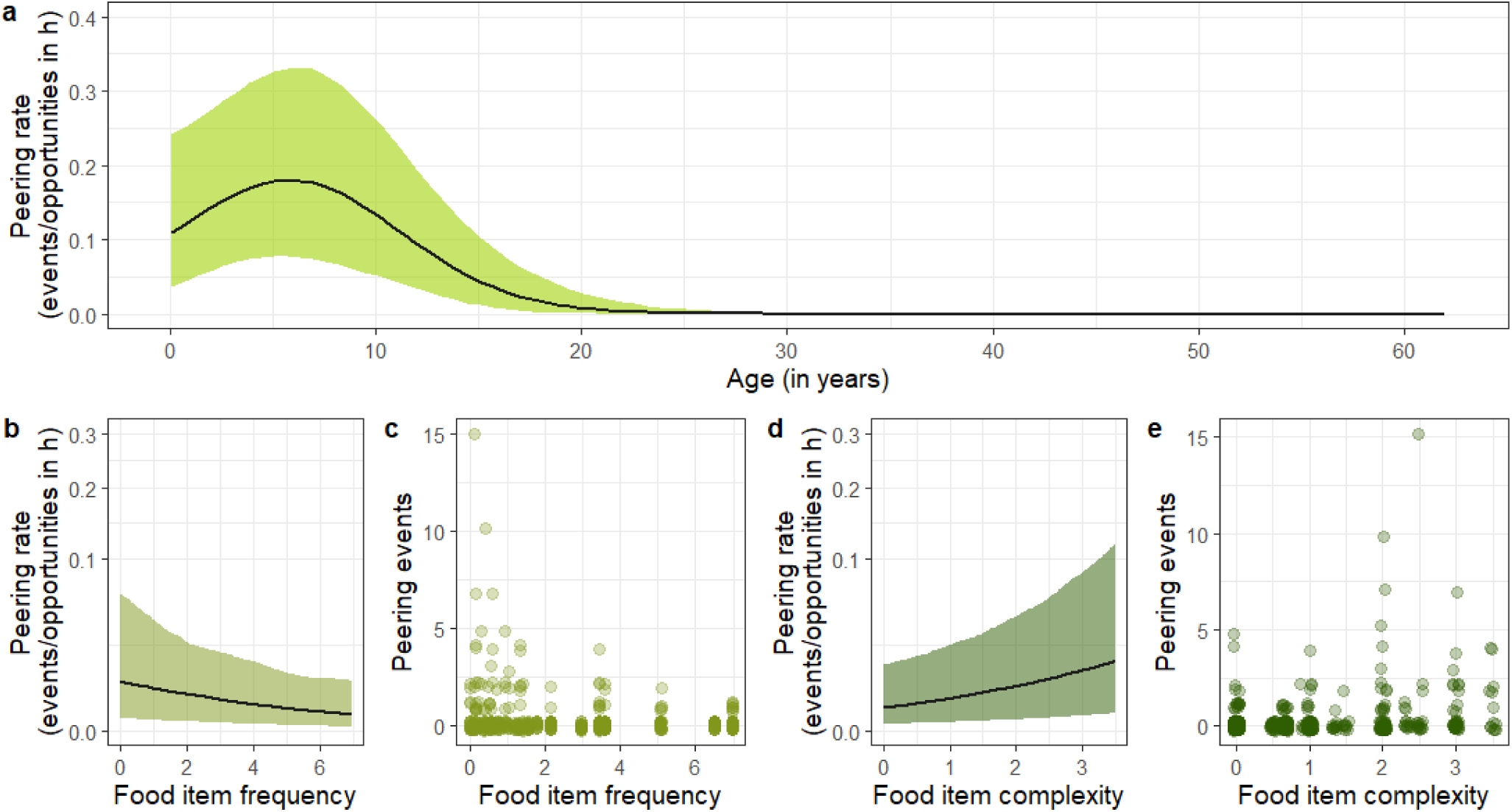
Effects of focal age, food item frequency, and food item complexity on peering (predictions 1a-b, 2c). Predicted peering rates a) per food item over age (yellow-green), b) over food item complexity (light olive) and d) frequency (dark green); black lines represent mean model predictions with 95% credible intervals as shaded areas; b) and d) are shown with square-root-transformed y-axis; number of observed peering events per c) food item frequency and e) complexity level; circles represent number of peering events of a given follow.

### Model B: Effect of target age on peering rates across development (prediction 1c)

Across age, chimpanzees showed significantly different peering rates targeting older, similarly aged, and younger conspecifics (full-reduced model comparison: LOO-CV ELPD=-33.0, SD=8.2). In particular, focal individuals peered at older conspecifics more frequently than at younger conspecifics throughout development, peaking at 4.9 years of age, Figure 3. Moreover, immatures peered at similarly aged conspecifics only at around 7.4 years of age, Figure 3b). The overall full model explained 37% [27%, 48%] of variation in peering rates. See Table S4 for model output and Figure S4 for HDIs.

**Figure 3:**
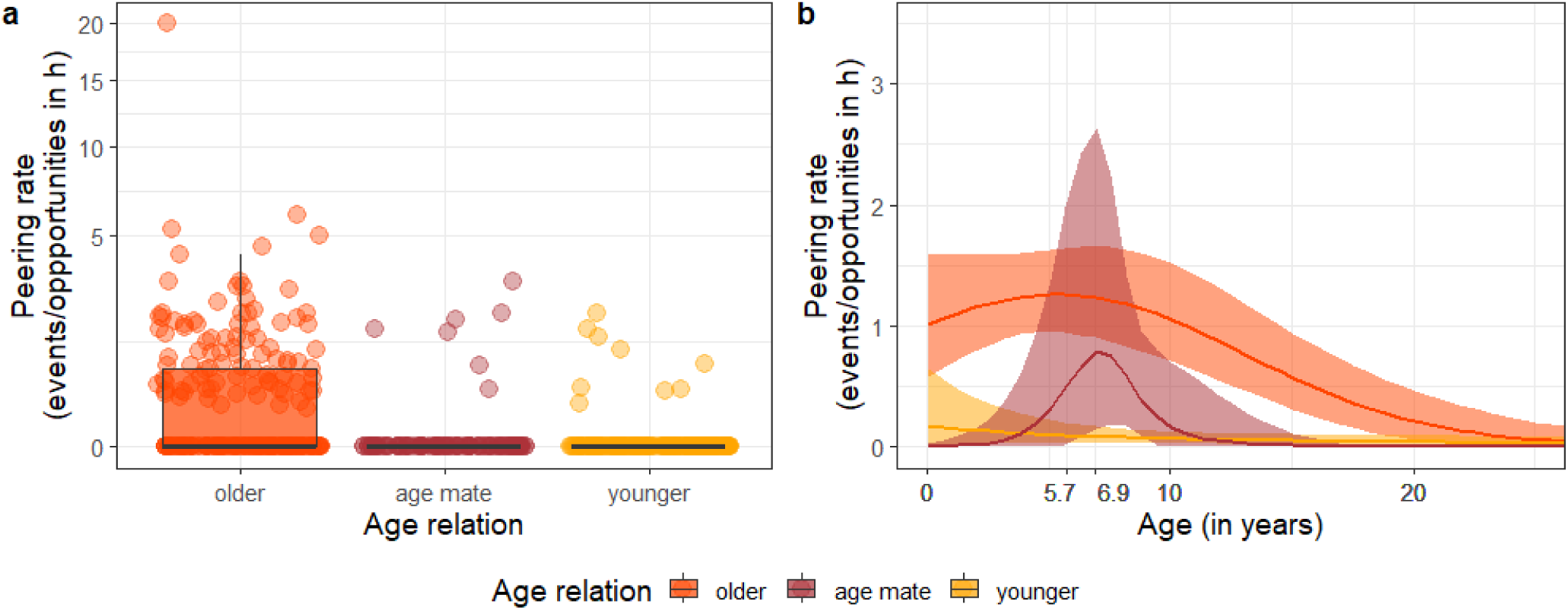
Effect of target age on peering rates across development (prediction 1c). a) Peering rates with respect to the age relationship of the peerer to the target (bold black line represents the median, box represents the 25%-75% quartile range, whiskers represent 1.5*interquartile range, points represent observations per follow per category), shown with square-root-transformed x-axis; b) predicted peering rates over age (only ages 0-25 are shown) by age relation category (older in orange, age mate in dark red, and younger in yellow); bold lines represent mean model predictions with 95% credible intervals as shaded areas.

### Model C1: Effect of kin relation on overall peering rates (prediction 2d)

Chimpanzees showed age-dependent differences in peering rates at targets of different kin relatedness, as indicated by a significant full versus reduced model comparison (LOO CV ELPD=-34.9, SD=7.8). In particular, mothers received most attention from the youngest infants. While this attention to mothers decreased with the immature’s age, mothers remained the most frequent peering targets until immatures reach around 5.9 years of age, at which point unrelated conspecifics became preferred peering targets, Figure 4a. Accordingly, focal individuals of average age (8.8 years) peered most frequently at unrelated individuals as compared to the mother (estimate=-1.15 [-1.94; - 0.39]) and other maternal kin (estimate=-2.76 [-4.26, -1.55]). The full model accounted for 46% [26%,62%] of variation in peering rates. See Table S5 for model output and Figure S5 for HDIs.

**Figure 4:**
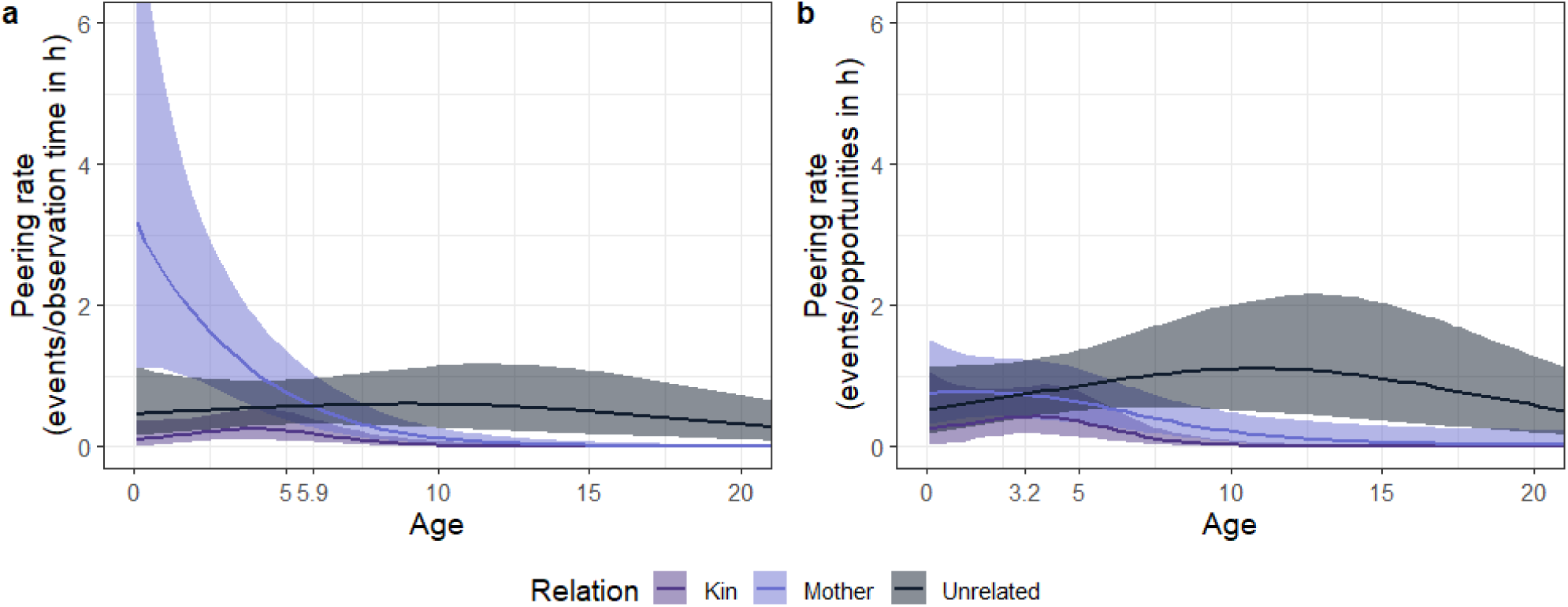
Effect of kin relation on peering (prediction 2d). Peering rates with respect to the kin relationship between peerer and target (kin in purple, mother in light blue, and unrelated in grey) over a) observation hours and b) peering opportunities (in hours); bold lines represent mean model predictions with 95% credible intervals as shaded areas.

### Model C2: Effect of kin relation on peering opportunity use (prediction 2d)

Peering rates (i.e. peering frequency controlled for opportunities to peer) targeting individuals of different kin relatedness (mother/kin/other) differed significantly over age (full-reduced model comparison: LOO CV ELPD=-24.2, SD=6.8). Focal individuals of average age (7.7 years) predominantly peered at unrelated individuals, as compared to mothers (estimate=-1.00 [-1.64, -0.39]) and other maternal kin (estimate=-2.37 [-3.43, -1.40]). Young individuals made early and increasing use of opportunities for peering at unrelated individuals, resulting in a shift of the primary peering targets from their mothers to unrelated individuals at around 3.2 years of age, Figure 4b. Effect size showed that the full model explained 58% [38%, 73%] of the variation in peering rates. See Table S6 for model output and Figure S6 for HDIs.

### Model D: Effect of conspecific dominance rank on peering frequencies (prediction 3b)

Chimpanzees did not show differences in peering rates targeting high, medium, or low-ranking female and male conspecifics, as revealed by the full-reduced model comparison (LOO-CV ELPD=-0.9, SD=3.3), Figure S7. For model output see Table S7 and for HDIs Figure S8.

### Model E: Increased daily grooming rates on peering days (prediction 3c)

Daily grooming rates for individual dyads were higher on days when one dyad partner peered at the grooming behaviour of the other dyad partner, as evident from the full-reduced model comparison (LOO CV ELPD=-8.6, SD=2.9, Figure 5). The full model revealed that this effect was driven by the binary peering predictor in the zero-inflated part of the model (estimate: -14.00 [-41.84; -2.93]): The probability of two chimpanzees to generally engage in grooming on a given day is strongly predicted by the presence of peering. On the other hand, once two chimpanzees engage in grooming, their time spent grooming does not differ between days when peering did or did not take place (estimate: 0.00 [-0.17; 0.17]). The full model explained 46% [41%; 50%] of variation in grooming rates. For model output and HDIs see Table S8 and Figure S9.

**Figure 5:**
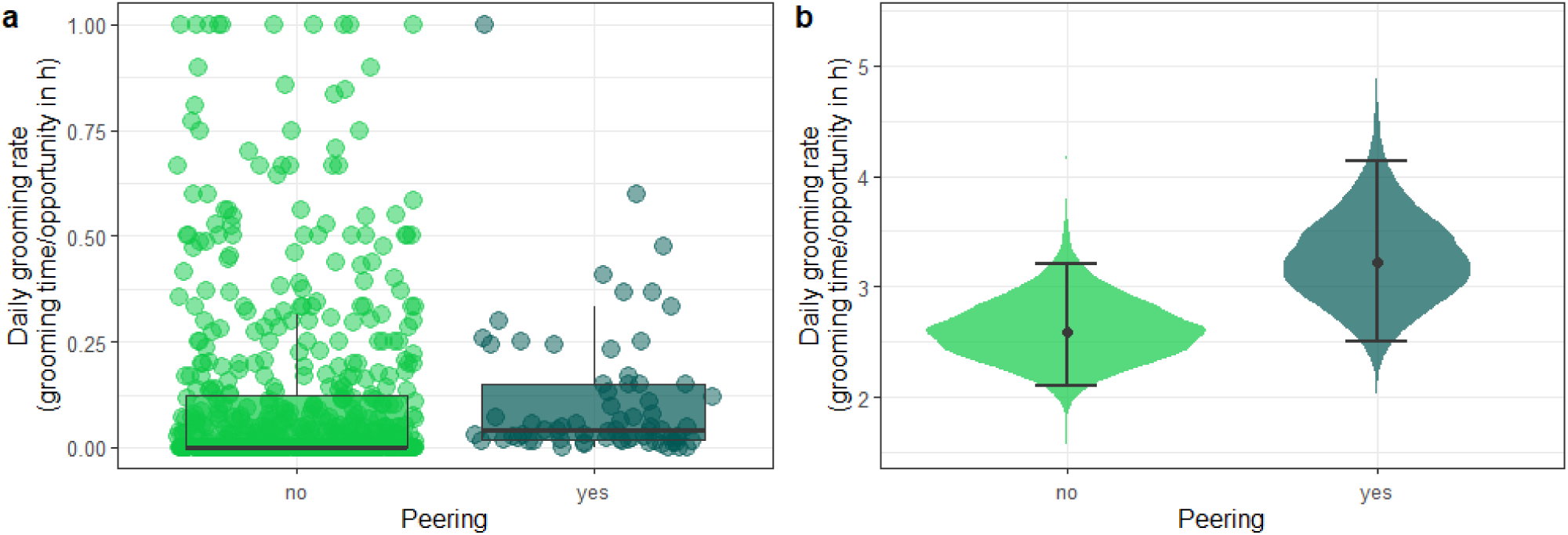
Increased daily grooming rates on peering days (prediction 3c). Daily grooming rates on days without and with peering (in light green and dark green respectively). a) Relative daily grooming rates by daily follow and dyad (bold black line represents the median, box represents the 25%-75% quartile range, whiskers represent 1.5*interquartile range, points represent observations per dyad and follow per category). b) Model predictions for relative daily grooming rates per category (points show estimates of conditional effects, error bars represent 95% credible intervals, violin shape displaying density distributions of estimates).

### Peering across the behavioral repertoire (predictions 2a-b)

To assess the range of peered-at behaviors relative to the behavioral repertoire of the Sonso community, we generated a full behavioral repertoire of all focal individuals and close party members (<5m) from scan sampling observations. Overall, we found that the observed behavioral repertoire consisted of 166 distinct behaviors of the categories Feeding (*n*=100), Social (*n*=36), Object Use (*n*=8), Exploration (*n*=7), Grooming (*n*=4), and other behaviors (*n*=11), see Table S9. This repertoire included 4 peering target behaviors that were never recorded during scan sampling and additionally 30 behaviors that were recorded for focal individuals but not for close party members, suggesting highly infrequent opportunities for the focal individual to peer. Focal individuals peered at 69 distinct behaviors across all behavioral contexts (Feeding *n*=41, Social *n*=14, Object Use *n*=4, Exploration *n*=4, Grooming *n* =3, Other *n*=3), Figure 6. The majority of peering (79%) was directed at feeding behaviors (*n*=187, including 9 feeding events on unidentified species) and grooming (*n*=92), followed by peering at social interactions (*n*=30), object use (*n*=20), exploration (*n*=17), and other behaviors (*n*=7).

**Figure 6:**
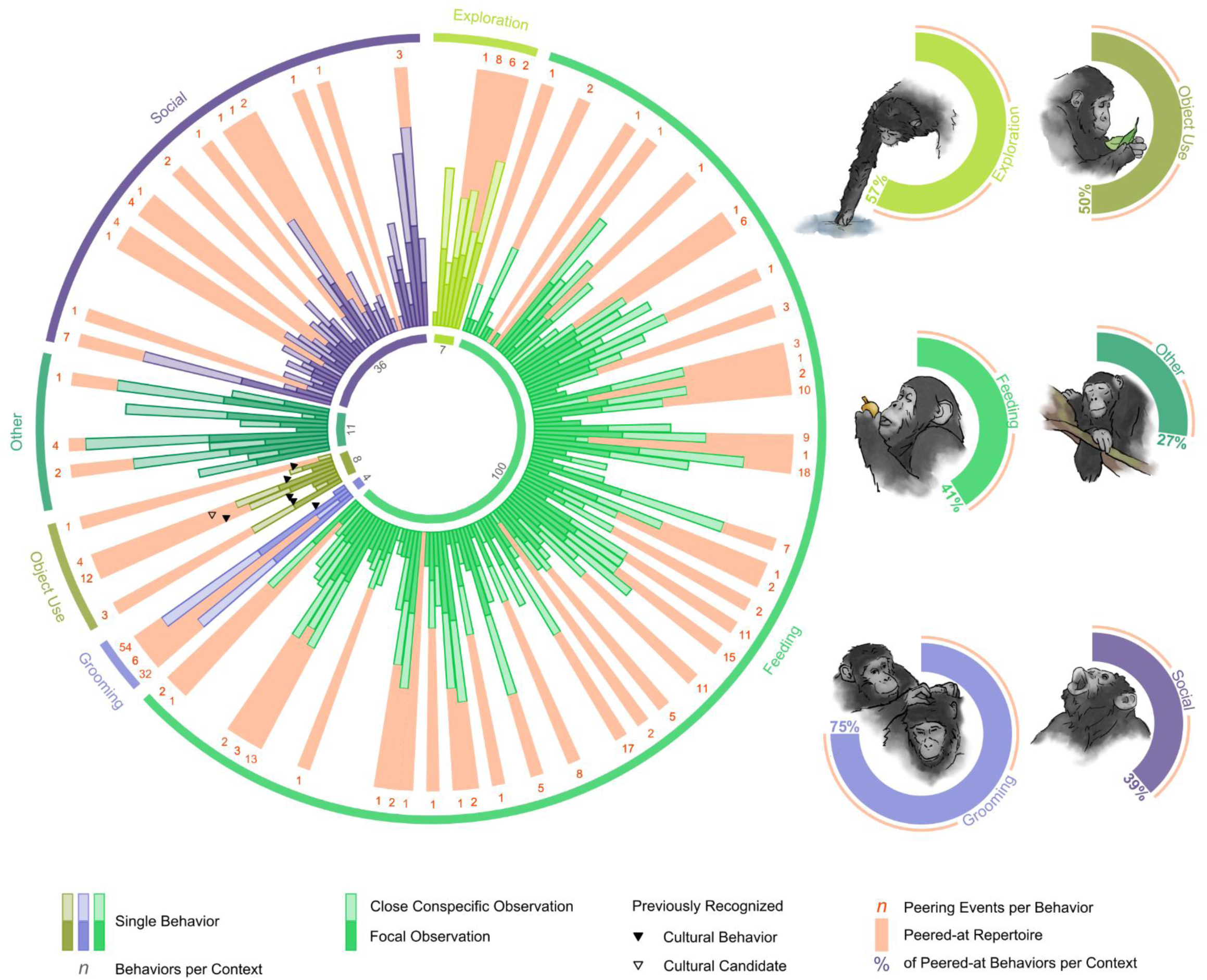
Peering across the behavioral repertoire (predictions 2a-b). Left: Bars represent log-transformed occurrences (original range *n*=0-5237) of unique behaviors by class of individual (occurrences of focal individuals in solid hue, of close party members in transparent hue) and behavioral context (Exploration in light green, Feeding in green, Grooming in light purple, Object Use in olive, Other in dark green, Social in dark purple). Black triangles show previously identified cultural behaviors (solid; from top to bottom: leaf-dab, ectoparasite inspection/squash on leaf, leaf-clip, seat-vegetation, handclasp-grooming) and candidates (outline; leaf-sponge). Red bars indicate peered-at behaviors with number of peering events per behavior (red numbers). Right: Proportion of peered-at behaviors by context with proportion percentages (same colors as left).

#### Additional peering functions (predictions 3a-c)

Lastly, we looked at potential additional functions of peering, namely peering as a 1) begging gesture, 2) a submissive signal, and 3) a signal to initiate affinitive interactions. 1) 235 out of 366 peering and active begging events were directed at behaviors involving items (e.g. fruits, roots, wood). Item transfer rates did not differ between active begging alone (62.5%) and combination of active begging and peering (69.6%; Fisher’s exact test: *p*=0.70). However, item transfer rates differed significantly between peering events (0%) and events which involved active begging (68.5%; Chi-Square Test of Independence: *X*^2^=145.36, *p*<0.00), Figure 7a. 2). We found that adult individuals peered at lower ranking conspecifics as frequently as at higher ranking ones (lower *n*=8, higher *n*=9; Exact Binomial Test: *p*=1), when comparing precise individual rank and assuming immatures to be lower ranking than adults, Figure 7b. 3). We found that 13% of peering was followed by social grooming within five minutes of the event (total *n*=312, grooming *n*=41) and rates differed significantly between contexts (Fisher’s exact test: *p*<0.000), Figure 7c. For peering in social contexts (total *n*=124), 29% of instances were followed by the peerer and target jointly engaging in the target activity. This affinitive interaction only happened following self-grooming, ectoparasite inspection with leaves, social grooming, and play, Figure 7d.

**Figure 7:**
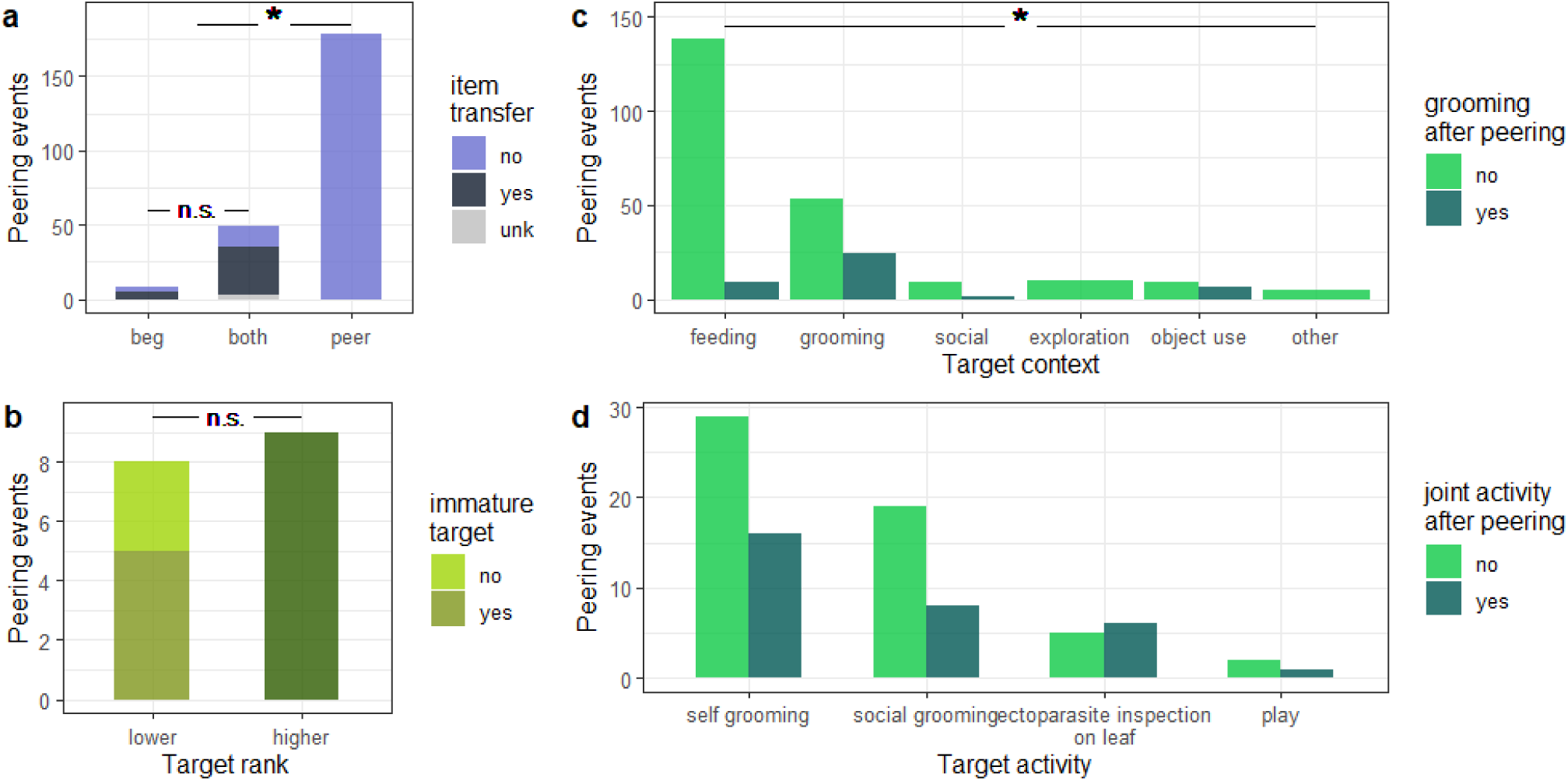
Additional peering functions (predictions 3a-c). a) Number of peering, active begging, and combinatory events that resulted in item transfer (item transfer in black, absence in blue, unknown in grey); b) number of peering events directed at higher and lower ranking individuals (immatures in light olive, lower ranking individuals in yellow-green, higher ranking individuals in dark olive); c) number of peering events per behavioral context that were followed by peerer and target grooming (grooming in dark green, absence in light green); d) number of peering events in social contexts followed by peerer and target engaging in the peered-at activity (joint activity in dark green, absence in light green); black lines represent comparisons across groups (n.s. showing non-significant and * significant results).

## Discussion

Decades of research on the spread of cultural elements in chimpanzees have highlighted the species’ substantial and sophisticated cultural repertoires^10^. However, the breadth of chimpanzee culture is less well-researched and, thus far, our knowledge was likely biased by methods focusing on showing the presence of cultural traits, rather than their full extent^40,78^. Here, we investigated evidence for observational social learning in chimpanzees to improve our understanding of the breadth of non-human primate culture by first validating peering as a behavioral indicator of social information seeking, then mapping the contexts in which social information seeking happens, and lastly, investigating potential additional functions of peering.

We found several lines of evidence that peering is a behavioral indicator of social information seeking in chimpanzees: following our predictions (1a-c) and studies on other primates^56,59^, peering rates peaked during immaturity, especially for infants and juveniles, while we found only a little peering in young adults and near zero rates of peering later in life. Peering therefore fell within the developmental timeframe in which we expect most learning to take place, as has been shown in orangutans^56^ and capuchins^59^. In line with previous research on chimpanzees^63^ and other non-human primates^56,73^, individuals further showed increased attention towards more complex and less frequently encountered food items, demonstrating that individuals peered selectively in learning-intense contexts. Peering was also generally directed at older – and to a lesser extent similarly aged – conspecifics, demonstrating that chimpanzees in this study (similar to western chimpanzees^63^, orangutans^79^ and capuchins^59^) selectively chose more experienced targets. Overall, chimpanzees peered in ways that suggest targeted social information seeking is likely to result in learning.

Our results align with previous research, which has demonstrated the importance of observational social learning for the acquisition of complex cultural skills in chimpanzees^18–21,25,26,30,31,63^. Not only did more complex feeding techniques elicit higher peering frequencies, individuals paid heightened social attention to tool and object use: Despite the fact that over the course of our study, juvenile and adult focal individuals engaged in tool or object use on average only once per daily follow, often for only a few seconds, 17 of the total 358 peering events in our data were directed at tool and object use. In line with this focus on complex but rare skills, we further found that rare food items induced increased peering (Figure 3) and that individuals made frequent use of opportunities to peer at uncommon behaviors such as wound treatment^80^. This selective use of infrequent learning opportunities could promote transmission of cultural skills, many of which occur only rarely. In this way, our results support the idea of observational social learning as a vital mechanism of cultural transmission and to corroborate previously identified cultural traits^10^.

However, in line with our prediction (2c), our results suggest that in chimpanzees, social information-seeking is not limited to complex behaviors like tool use. Instead, peering events targeting tool or object use constituted only a small proportion of the overall number of peering events in our dataset (Figure 6). Most of the peering (∼80%) was directed at (non-tool) feeding and grooming behaviors, and therefore at less conspicuous, everyday activities. Acquiring proficient feeding skills is crucial for immatures as they gain independence from their mothers and need to independently sustain their energetic needs^68^. It is therefore not surprising that young chimpanzees make use of social learning opportunities to acquire a broad range of adult feeding skills. Importantly, while we have shown that complex feeding techniques and rare food items received greater attention relative to their occurrence, we found peering behavior targeting all levels of food item complexity and rarity (Figure 2). This widespread occurrence suggests that immature chimpanzees use observational social learning to acquire substantial shares of their feeding repertoires. High peering rates at feeding behavior compared to other behavioral contexts further highlight the potential for diet repertoire and feeding techniques to encompass extensive cultural knowledge. The cultural relevance of simple feeding skills is easily missed by conventional group-level approaches of mapping cultural traits (e.g. the method of exclusion^10,40,78^), as socially acquired feeding competencies remain invisible to group comparisons for several reasons: On the one hand, socially acquired diet repertoires of populations inhabiting similar habitats might show high degrees of overlap because of similarities in food availability. Similarly, feeding techniques might follow similar behavioral sequences because of overlapping food item properties and limited variability in functional pre-ingestive processing solutions. On the other hand, where socially transmitted feeding skills vary alongside ecological differences, they may go unrecognized. Lastly, socially transmitted feeding skills may go unrecognized when they are not studied in sufficient detail to detect subtle variation. For example, variation in termite fishing was found in the resting posture chimpanzees adopted while fishing.

Besides frequent peering at feeding, chimpanzees often observed conspecifics grooming. While grooming constitutes a primate universal and thus likely emerges through hard-wired development, the high peering rates at grooming suggest that specific elements of the behavior may be socially learned. Previous research has shown that certain grooming patterns in chimpanzees can be ascribed to cultural knowledge, such as handclasp grooming^45,81^. Previous research has also shown more fine-grained group differences in grooming techniques: Ngogo chimpanzees in Uganda were observed using a poking motion, while Mahale chimpanzees in Tanzania use a stroking motion to scratch their grooming partner^82^. Grooming techniques seem to differ in similar ways between the Sonso and the neighboring Waibira chimpanzee community (personal observation C.H.). Furthermore, in Japanese macaques (*Macaca fuscata*) grooming style has been shown to be transmitted socially^83^. Overall, our results add to previous findings suggesting that the development of grooming techniques is mediated through social knowledge.

Chimpanzee cultures have previously been estimated to comprise a total of around 39 behavioral variants across populations, identified from a selection of 65 putative cultural traits^44^. In contrast, we recorded a minimum of 69 unique peering target behaviors in a single chimpanzee community— most of them (*n*=41) being (non-tool) feeding behaviors. Only two of these behaviors (leaf-dab, ectoparasite inspection/squash with leaf^44^) had previously been recognized as cultural. The 69 unique target behaviors in this study do not comprise potential variants of behaviors (e.g. different feeding techniques for one food item, different grooming styles, different nesting elements, or different object manipulations). As previous research has demonstrated cultural variation in expressions of individual behaviors^13,33,38,50,82^, our approach of counting unique peering target behaviors but not variation in their expression represents a highly conservative estimate of cultural candidates for the Sonso chimpanzee community. This number of cultural candidates nonetheless almost doubles the previous estimate of cultural behaviors in chimpanzees. Assessing cultural transmission by studying behavioral indicators of social learning, thus suggests a wide, unrecognized range of potentially cultural traits in chimpanzees and can substantially contribute to a better understanding of the true breadth of non-human primate cultures.

Social learning has been proposed to follow three distinct phases characterized by differences in target choice^72^. Our results support the first two phases of learning: up until ∼5.9 years of age, immature chimpanzees peered most frequently at their mothers (Phase 1), and afterwards increasingly targeted other group members (Phase 2). Heightened attention towards the mother closely follows the average duration of inter-birth intervals at Budongo^84^, suggesting that changes in peering target choice might result from adjustments in the mother-offspring relationship^85^. However, when taking the availability of role models into account, we found that young chimpanzees leveraged opportunities to peer at unrelated group members from an early age (Figure 4b). In fact, we found that mothers receive only slightly more attention relative to unrelated conspecifics and for fewer years (∼3.2 years, Figure 4). Recent research on peering in chimpanzees interpreted high peering rates directed at mothers as a result of their high social tolerance toward peering immatures^63^. However, accounting for opportunities to peer, our results suggest that the focus on mothers as role models is likely explained in part by them being more available to immatures, as compared to other conspecifics. Maternal kin did not constitute frequent peering targets, neither in absolute terms, nor when taking peering opportunities into account. However, our data set only comprised limited data on opportunities to peer at maternal kin, allowing for limited conclusions only. The subsequent focus on unrelated group members comprised a period of heightened attention towards similarly aged conspecifics: male juveniles of ∼7 years of age (n=4, no female in this age range was present in the group) closely observed their peers (Figure 3b). As juvenile chimpanzees of this age gain more independence and engage more frequently with a broader range of group members^86^, increased attention to other juveniles might – particularly in philopatric males^86^ – stem from wanting to join and conform to their peer group^71,87^. However, three of these four juveniles were orphans (two without any maternal kin), limiting the generalizability of our conclusions. Overall, these patterns on peering target selection suggest that immature chimpanzees use learning opportunities that go hand in hand with the species’ high levels of sociability: they peered at a variety of targets from an early age and continued peering at a variety of group members throughout immaturity.

The fact that peering continued throughout immaturity could point to the need for a prolonged period to acquire and master skills, and thereby support the needing-to-learn-hypothesis^88,89^. Notably, we found particularly high peering rates for juvenile chimpanzees in feeding contexts (Figure 2). Previous research on feeding skill acquisition in chimpanzees^90^, as well as in other primates^91–96^, has shown that immatures reach adult-like feeding skills, such as food recognition and processing skills around weaning. This threshold has been interpreted to demonstrate that immatures do not need to learn feeding skills after nutritional independence. However, learning into adulthood has been identified in complex skill acquisition, with previous research on tool use^97^ and feeding^63^ showing that the acquisition of cognitively challenging tasks takes the most time. In our study, juvenile peering in feeding contexts could not be ascribed to heightened attention to complex or rare – potentially more cognitively demanding – feeding techniques (Table S3), but we found a trend that peering at more complex food items peaked later in immaturity (Figure S3). No or only a limited influence of feeding complexity on prolonged peering behavior indicates that chimpanzees might improve general aspects of their foraging and feeding skills after nutritional independence, such as increasing their diet repertoire or their knowledge on feeding locations.

As with many behaviors, the immediate motivation to engage in the behavior may differ from its evolved function. Furthermore, behaviors may have more than one evolved function. Even though our results highlight the use of peering in learning-relevant contexts, peering behavior is unlikely to be elicited by a conscious desire to learn or seek information. The underlying motivations might vary across contexts and as a function of age or a combination thereof. We found that rare behaviors elicited the highest peering rates, suggesting that at the immediate level, curiosity, i.e. a want to understand and an inclination towards novelty^98–101^, may drive peering behavior. In line with previous research on begging in chimpanzees^63^, our results show that it is unlikely that peering evolved as a form of begging (i.e. to obtain items). However, the motivation to get an item may, at times, nonetheless elicit peering. This may be especially true for high-value food items such as meat, where active begging likely requires higher levels of social tolerance and may carry a greater risk of retribution than peering. Peering could further constitute a way to assess social tolerance and initiate active begging accordingly. However, the small number of combinatory events (peering with active begging) renders this unlikely. We further assessed potential social functions of peering, such as signaling submissiveness to more dominant individuals and initiating affinitive interactions. Our results did not show an effect of association partners’ dominance ranks on peering frequencies (Figure S7), nor did we find that peering unilaterally followed the dominance hierarchy of adult individuals (Figure 7). This pattern indicates that peering was neither motivated by interest in high-ranking conspecifics nor did it function as a submissive signal. Instead, our data suggests, that depending on social contexts, peering might function as a signal of interest or willingness to initiate affinitive interactions: we found that peering at social and self-grooming was occasionally (∼34%) followed by grooming between the peerer and the peering target, and particularly peering at ectoparasite inspection on leaves – while rare (n=11) – mostly (55%) resulted in joint social grooming (Figure 7). In addition, we found that dyads groomed relatively more on days when a dyad partner peered at grooming behaviors of the other partner, suggesting that peering can signal interest to engage in social grooming (Figure 5). Matsuzawa^71^ and de Waal^87^ proposed that paying close social attention to group members follows from an immature’s want to belong and to conform to the group. Signaling initiative to engage in affinitive interactions, frequent peering at mothers, as well as the juveniles’ heightened interest in age-mates in our study, are all in line with this hypothesis. A want to belong and conform may thus constitute an important motivation to peer. Overall, while peering is likely driven by various immediate motivations and might hold multiple adaptive functions, the attentive observation of conspecifics enables knowledge transfer and thus observational social learning. Diverse underlying motivations and additional functions could drive the emergence of cultural variants across a broad range of contexts.

In conclusion, our results suggest that peering is a suitable behavioral indicator of targeted social information-seeking and a likely means of observational learning in wild chimpanzees. Immature chimpanzees peered at a wide range of behaviors across contexts, including simple everyday skills such as (non-tool-assisted) feeding and grooming. Chimpanzee everyday behavioral repertoires – beyond conspicuous, complex skills – thus likely encompass a broad, previously underappreciated body of cultural knowledge. We found mothers to be frequent peering targets of infant chimpanzees. However, chimpanzees leveraged opportunities to peer at unrelated conspecifics from a young age and predominantly peered at unrelated conspecifics during later development. Hence, while mothers likely constitute important cultural vectors, the high sociability of chimpanzees may foster diverse pathways of cultural transmission. Lastly, although we did not find evidence that peering functions as a begging or submissive gesture, our results suggest that peering could signal attempts to initiate affinitive interactions in social contexts. Peering might be driven by various immediate motivations, such as curiosity, a want to obtain an item, or a wish to belong and conform. This diversity in functions and motivations likely stimulates cultural transmission of a broad variety of behaviors.

## Methods

### Ethics

This research was approved by the Budongo Conservation Field Station, the Ugandan Wildlife Authority (UWA, reference number: COD/96/05), and the Ugandan National Council for Science and Technology (UNCST, reference number: NS319ES). The study is based on observational data collection on a well-habituated chimpanzee community, and observational data were collected in adherence to the International Primatological Society’s Code of Best Practice for Field Primatology^102^.

### Study Population

The Sonso community in the Budongo Forest, Uganda, (01u43’N, 31u32’E) has been studied since 1990 and from 1994 most male chimpanzees were habituated to human observers^84^. Throughout the study period, the community consisted of 70-75 individuals. For this study, we collected data on 28 individuals, 11 adults (5 females) and 17 immatures (7 females) ranging from 1.5 months up to 62 years of age. Ages were classified as adults (15+ years) and immatures: infants (0-5years), juvenile (5-10 years), subadults (10-15 years)^84^. Ages of individuals born prior to the establishment of the field site had been estimated based on inter- and intra-community comparisons of visual features with individuals of known age^84^. Similarly, ages of immigrating females were estimated based on visual features.

### Data Collection

Data were collected from January 2022 to August 2024 during focal follows. Focal individuals were followed between 7:00 am and 4:30 pm; whereas the active period of the chimpanzees usually lasts from ∼6:30 am to 6:30 pm^84^. As a result, the observation periods did not comprise all daily activities (e.g. construction of evening nests). We collected data via instantaneous scan sampling at 3-minute intervals. At each scan, we recorded the activity of the focal individual and all individuals within 5 meters of it. Whenever the focal individual was feeding, we recorded the feeding species and ingested part (e.g. plant parts including fruits, leaves, flowers, bark, pith, wood, insects and insect nests, meat), which when combined are henceforth referred to as “food item”^91^. The data collection protocol was adapted from the standard orangutan data collection protocol of the orangutan network^103^. In addition to scan sampling, we collected all-occurrence data on peering and active begging by the focal individual (see Table 1).

**Table 1:** Definitions of terms used in this study.

|  | Definition |
| --- | --- |
| Peering | Close-range attentive observation of a conspecific (target/role model) for more than three seconds. Distance to target must be less than 5m and allow detailed observation of the behavior (Figure 1). |
| Active Begging | Active reaching with hand or mouth towards an item (food item or other object, e.g. leaf, stick, shell) in possession of a conspecific (target). Can involve an active attempt to gain access to the, either by attempting to or grabbing the item or by attempting to or taking the item in the mouth. |
| Item Transfer | An item is obtained from a conspecific, e.g. through scrounging or begging. |
| Affinitive Interaction | Joint activity of two or more individuals, e.g. social grooming, play. |

In total, we collected 1136h of observation hours with a mean observation time of 40.6h per focal individual (min=14.8h, max=56.5h, median=43.3h) and a total of 366 peering events (including 8 active begging events and 49 combinatory events of begging and peering). Data were collected by N.E.S., M.G.M., A.S.P., and R.Y. All observers reached a minimum of 87% agreement with N.E.S. on scan sampling data collected during simultaneous follows before their data were included in the dataset. Peering data were too intermittently observed to include in a quantified inter-observer assessment, but data collection on peering was thoroughly discussed and monitored by N.E.S during several weeks of training observations.

All behavior of conspecifics within 5m of the focal individual were considered potential learning opportunities for the focal individual. For each conspecific, kin relations to the focal were categorized as “mother”, “kin”, and “unrelated”, where “kin” includes all maternal kin (except the mother) up until third-degree relatedness (e.g. cousins). Young chimpanzees live in maternal family units and are frequently exposed to maternal kin^104^, whereas paternal care and likely also the recognition of paternal kin is greatly limited^105,106^. Age relations between the focal and conspecifics were defined as similarly aged “age mates” for individuals within +/-1.5 years of the focal, younger, and older individuals.

We used long-term field data collected by local field assistants over 6 years of focal follows (2018-2023) to assess the frequency of food items in the population’s diet. We expressed the frequency of each food item as a percentage of total population feeding time, i.e. the summed hours all independent individuals spent feeding on an item divided by the total recorded feeing hours. One food item (fruit of *Ficus mucuso*) made up over 25% of the observed diet, with all other items ranging between 0.1% and 6%. Because the prevalence caused a highly uneven data distribution, *Ficus mucuso* was excluded from the analysis. Data on food processing complexity were compiled by N.E.S. and G.M., based on feeding videos and personal observations. In accordance with previous studies^91^, for each food item, pre-ingestive processing steps (e.g. “bite open”, “extract flesh”, “spit out seeds”) were listed and scored, and where necessary (i.e. where several techniques for the same food item were present) the score was averaged over multiple feeding variants.

Chimpanzee groups follow a largely linear dominance hierarchy, with males generally ranking higher than females^104,107^. Long-term field data on submissive greetings, “pantgrunts”, were used to calculate hierarchical rank as Elo-Ratings ^108^ using the R package “EloRating”^109^. Rank was derived from pantgrunt data from 2017-2024 for females and data from 2021-2024 for males. Male rank was calculated per month of the study period, female rank was calculated overall. Rank was only calculated for chimpanzees 14 years or older, leading to a hierarchy of 41 ranks (10 male, 31 female). Individuals were sorted into “high”, “medium”, and “low” rank classes per sex, where each category comprises standardized elo-ratings higher than 0.5, between 0.3 and 0.5, and lower than 0.3 respectively, with the alpha male being included in the class “high male”. Females were ranked below males, and all immatures were assumed to be lower ranking than ranked individuals.

### Analyses

Descriptive analysis: Peering across the behavioral repertoire (predictions 2a-b)

We generated a behavioral repertoire of the Sonso community for our study period by compiling all unique behaviors observed during scan sampling of focal individuals and conspecifics within 5m of the focal. Classifying When close conspecifics were never seen to engage in a behavior, peering opportunities for the focal individual were likely highly infrequent. We categorized these behaviors by their context and checked whether they had previously been recognized as cultural traits of the Sonso community^44^. This behavioral repertoire was matched with the peering data, creating additional behaviors where scan sampling had not captured a peered at behavior. For all unique behaviors, the number of occurrences during scan sampling (focal/party member) and the number of peering events were counted. The resulting behavioral repertoire and set of unique peering target behaviors did not include different behavioral variations (e.g. different feeding techniques for the same food item, different grooming techniques).

Statistical tests: Additional peering functions (predictions 3a-c)

We investigated 1) whether peering serves as a begging gesture in wild chimpanzees by assessing how frequently peering and active begging occurred separately and jointly, and how frequently these occurrences resulted in item transfer, based on 366 peering and active begging events of 25 individuals. We ran Fisher’s Exact Test in R (version 4.3.3^110^) using function fisher.test() to compare item transfer rates following active begging alone and combinatory events of active begging and peering. We then assessed differences in item transfer rates between sole peering events and events that involved begging with a Chi-Square Test of Independence using function chisq.test().

Further, to investigate whether chimpanzees used peering to signal submission, we assessed whether peering at lower- or higher-ranking individuals differed from chance (p=0.5) using an Exact Binomial Test, function binom.test(). This analysis was only based on adult peerers and their targets, because immature individuals were not included in rank categorization but were assumed to be generally lower ranking than adults. This analysis was therefore based on 17 peering events of 8 adult individuals (see below for additional analysis on effects of rank).

Lastly, to investigate whether chimpanzees use peering to initiate affinitive interactions, we assessed how frequently peering was followed by social grooming between peerer and target. To identify whether frequencies of joint grooming after peering differed across behavioral contexts, we ran a Fisher’s Exact Test and visually inspected frequencies of joint grooming across contexts. We then looked at social contexts in more detail by assessing how often peering at social contexts (i.e. grooming, play) was followed by the peerer joining in the social situation, e.g. how often peering at grooming was followed by peerer and target grooming. Here, we included self-grooming and ectoparasite inspection on leaves as target contexts, as peering could signal the intention to join in the (self-) grooming behaviors (see also Model E for an additional analysis). The analyses on affinitive initiations were exclusively based on peering events for which respective information was available (265 peering events).

Generalized Linear Mixed Models: Peering as an indicator of observational social learning, peering target selection in relation to kin relatedness and individual rank (predictions 1a-c, 2c-d, 3b-c)

To assess the development of peering frequency over age and the effects of food item complexity, food item frequency, target kin relation, target age, and target rank on peering frequency, we carried out five separate statistical analyses: We fit five generalized linear mixed models (GLMMs) with a negative binomial error structure and logit link function using the Bayesian framework in R (version 4.3.3^110^), using Stan^111^ and the bmrs package^112^. For each model, we used the number of peering events as the response variable and offset the respective observation hours (see below) using the “rate” function of the bmrs package. In addition, to investigate whether peering has an effect on daily grooming rates, we fit a GLMM with a zero-inflated Poisson error structure and log link function, where the number of scans a dyad spent grooming was the response and the number of scans both dyad partners were present was the offset (implemented using the “offset” function). We fit all models with the brms’ default (uninformative) priors and ran them for four chains of 4000 iterations each, including 1000 warmup iterations. Adapt-delta was set to 0.95 to prevent divergent transitions. Model fitting resulted in effective sample sizes (ESS) ranging for bulk ESS from 958.34 to 14,663 and for tail ESS from 560.70 to 10,449.21 and Rhats of 1 for all estimated parameters, indicating good model convergence. We ruled out overdispersion for all models by visually assessing dispersion of observed and simulated data using functions “pp_check” and “ppc_stat” of package bayesplot^114^.

Because we predicted peering frequency to peak during early immaturity, Models A-D included focal age as a quadratic term (i.e. “age^2^”) besides linear age (“age”). All continuous predictors were z-transformed (mean of zero and standard deviation of one) to support model convergence and interpretation of model output. We included random intercept effects as controls (see below) to ensure the nominal rate of 5% for type I errors^115^. All five models were based on data of individuals for whom at least 20h of focal observations could be recorded. This comprises 27 individuals of ages 0.1 to 62 years with mean follow hours of 42.9h (min=20.4h, max=56.5h, median=45.1h).

A. Effects of focal age, food item frequency, and food item complexity on peering frequencies (predictions 1a-b, 2c): To investigate the development of peering frequency over age and the effects of food item complexity and frequency on peering rates, we fitted a model with “age”, “age^2^”, “item complexity”, and “item frequency” as predictors. To account for potential individual differences and effects of specific food items and to avoid pseudoreplication, we included “focal ID” and “item ID” as random intercepts. To account for varying opportunities to peer, the model was offset with log-transformed “opportunities”, i.e. duration (in hours) of conspecifics feeding on a specific food item during a given follow. The model is based on observations of 27 focal individuals (age range=0.1 – 62 years) over 1121 observation hours. See SI for a matched model including the interaction term of “item complexity” with “item frequency” (Model A.1) and a model including interactions of “item complexity” and “item frequency” with both age terms each (Model A.2). In this way, Model A.1 controlled for potential covarying effects of “complexity” and “frequency” (Figure S1) and Model A.2 assessed whether the effects of food item frequency or complexity changed with increasing age, i.e. with an individual’s feeding competence. Neither model showed significant effects of the interaction terms.
B. Effect of target age on peering frequencies across development (prediction 1c): To investigate how frequently individuals peer at targets of different ages relative to their own age, we fit a model with the predictors “age”, “age^2^”, and “age relation” (younger/age mate/older). To assess whether target selection based on target age classes changes throughout development, we further included the interaction of “age” and “age^2^” with the predictor “age relation”. The interaction term and the main effect “age relation” constituted the terms of interest. To further control for individual differences, we included “focal ID” as a random effect. This model included an offset term of opportunities describing the presence of at least one conspecific (in hours) of the respective age-relation category per follow. This analysis thus reflected relative peering rates for each target class. The model was based on 1113 hours of observation on 27 focal individuals (age range=0.1-62 years).
C. Effect of kin relation on 1) overall peering frequencies and 2) on peering opportunity use (prediction 2d): To investigate the amount of peering directed at individuals of different relatedness across an individual’s development, we built two models with identical fixed and random effects structure, namely the main effects and interaction of “age” and “age^2^” with the predictor “target relation” (mother/kin/unrelated). The interaction term constituted the predictor of interest, because we expected that relatedness to conspecifics would affect target selection differently across development. To account for individual variation and to avoid pseudoreplication on specific follow days, we included the random effects “focal ID” and “follow Nr”. The first model was offset with log-transformed “observation hours”, i.e. the observation duration. Thus, the first model did not take variation in peering opportunities into account but described general peering frequencies over age. The second model was offset with the opportunities to peer at the different role models, i.e. the duration (in hours) during which at least one individual of a certain relatedeness was present. Hence, the second model described peering frequencies relative to available peering opportunities. These models were based on 710 hours of observation on individuals for whom a mother was present in the community (*n*=18, age range=0.1-30.6 years).
D. Effect of conspecific dominance rank on peering frequencies (prediction 3b) To assess how frequently individuals peer at conspecifics of different rank, we ran a GLMM with predictors “age” and “age^2^”, as well as the interaction of and the main effects of “sex” (male/female) and “rank class” (high/medium/low). The interaction constituted the term of interest, because we expected high-ranking individuals (i.e. high-ranking males) to attract more social attention than lower ranking individuals (i.e. lower-ranking males, all females). To account for individual variation and avoid pseudoreplication on follow days, we included the random effects “focal ID” and “follow Nr”. The model was offset with opportunities to peer at different rank-sex classes, i.e. the duration (in hours) conspecifics of a specific rank-sex class were present throughout the follow. Opportunities and peering events were hence limited to the presence of adult (i.e. ranked) conspecifics. The model was based on 1089 observation hours on 28 focal individuals.
E. Increased daily grooming rates on peering days (prediction 3c) To assess if the presence of peering affects dyad-specific daily grooming rates, we ran a GLMM with the outcome “nr of grooming scans” and the log-transformed offset of “scans of partner presence”. To account for the high proportion of zeros in the data, i.e. for the high number of days on which dyad partners spent time in proximity but did not engage in grooming, we ran a zero inflated Poisson model. For both model parts, the modelled outcome “grooming scans” and the zero-inflated part, we included the predictor “peering” (yes/no) as the term of interest and the random intercept “dyad ID” to account for variation within specific dyads. The model was based on 692 observations on 27 focal individuals. We included only dyads who engaged in grooming at least once (145 individual dyads of focal and non-focal individuals).

We assessed model estimates and credible intervals (CD) to evaluate predictor effects, where we considered credible intervals not overlapping zero to strongly support the presence of a meaningful effect^117,118^. Additionally, for models B-E we ran full-reduced model comparisons to assess overall effects of categorical predictors and interaction terms. Reduced models had the identical model structure as full models, but lacked the predictor(s) of interest, and were compared using Leave-One-Out-Cross-Validation (LOO-CV)^119^ with function “loo” of the “loo” package^119^. This comparison estimated whether the full model resulted in an improved out-of-sample predictive performance by assessing the difference in the expected log predictive density (ELPD) of the full and the reduced model. We judged the full model, and hence the predictor(s) of interest, to hold additional explanatory power when the full model had the lower ELPD and the ΔELPD ± 2 × SE did not overlap zero, where ΔELPD refers to the difference between ELPDs of the full and reduced model and SE to the respective standard error. We report highest density intervals (HDI) of posterior distributions of model estimates (see SI Results section) and Bayesian R^2^, which we obtained using the “bayes_R2” function of the “bmrs” package.

## Resource availability

Requests for further information and resources should be directed to and will be answered by the lead contact, Nora E. Slania. Observational data and code are available at https://data.mendeley.com/preview/38v7jbvdw6?a=641fbf33-fe58-4164-8840-d4830d6d8fd5. The repository will be made public with submission.

## Acknowledgments

We are thankful to the Ugandan Wildlife Authority (UWA) and the Ugandan National Council for Science and Technology (UNCST) for permitting and supporting our research in the Budongo Forest. We are grateful to the staff at the Budongo Conservation Field Station (BCFS), management, field station staff, and field assistants alike for welcoming us and supporting us in and outside the forest throughout several years. We are especially grateful to the other Sonso field assistants, Chandia Bosco, Monday Mbotella Gideon, Adue Sam, Asua Jackson, as well as the site director, David Eryenyu, and research coordinator, Simon Peter Ogola. We thank Pascal Kuhn for his help with the figures and his work on the chimpanzee drawings. This study was funded by the Volkswagen Stiftung (Freigeist fellowship to C.S.) and the Max Planck Institute of Animal Behavior. T.R. was supported as a postdoc by the Marie Sklodowska-Curie Actions postdoctoral fellowship (project number: 101150646).

## Author contributions

Conceptualization: C.S., N.E.S

Data curation: N.E.S, C.H.

Formal analysis: N.E.S, T.R.

Funding acquisition: C.S.

Investigation: N.E.S., M.G.M., A.S.P., G.M., R.Y.

Methodology: C.S., N.E.S

Supervision: C.S., K.Z., N.E.S.

Visualization: N.E.S., T.R.

Writing – original draft: N.E.S., C.S.

Writing – review & editing: N.E.S., C.S., K.Z., C.H., M.G.M., A.S.P., G.M., R.Y., T. R.

## Supplemental Information

Supplementary Materials Document: Figures S1-S8, Table S1-S8

