## Supplementary Materials for "Chimpanzee culture beyond the conspicuous: Evidence for broad-scale observational social learning in wild individuals"

### Analysis

#### Statistical Analysis

Model A: Frequency Complexity Age

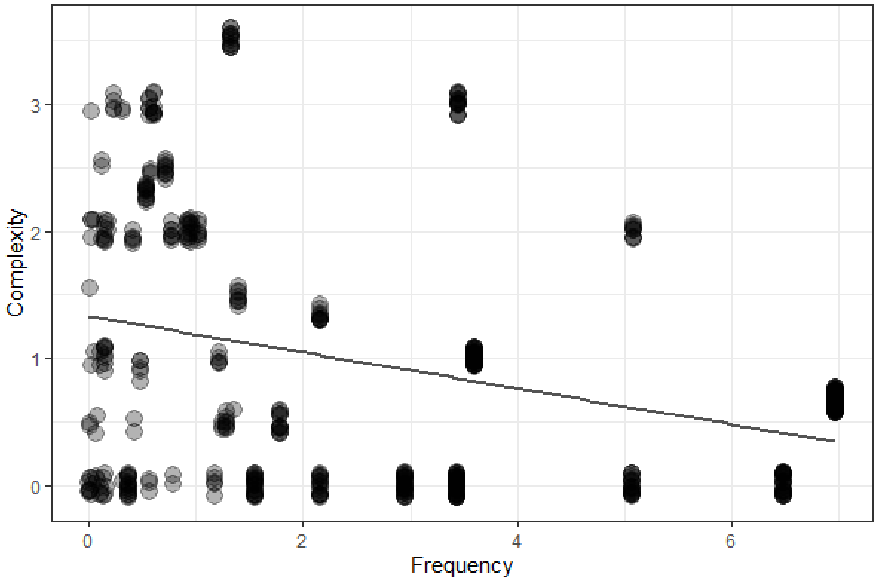


Figure S1: Distribution of food items by frequency and complexity, related to STAR Methods Model A.

### Results

**Model A:** Effects of focal age, food item frequency, and food item complexity on peering frequencies

Table S1: Model A output, related to Figure 2. Model estimates and respective standard error, credibility intervals (Q2.5-Q97.5), Rhats, Bulk ESS and Tail ESS. All predictors (“complexity”, “frequency”, “age”, “$age^2$”) were z-transformed. Credibility intervals not comprising zero are marked in bold.

| | Estimate | Est.Error | Q2.5 | Q97.5 | Rhat | Bulk_ESS | Tail_ESS |
|---|---|---|---|---|---|---|---|
| Intercept | -1.438 | 0.458 | **-2.394** | **-0.597** | 1 | 7276.509 | 8661.897 |
| comp_z | 0.642 | 0.209 | **0.249** | **1.075** | 1 | 5512.26 | 7039.836 |
| freq_z | -0.696 | 0.299 | **-1.269** | **-0.087** | 1 | 6062.381 | 7679.016 |
| age_z | -3.252 | 0.975 | **-5.374** | **-1.606** | 1 | 6066.579 | 6153.285 |
| age_z2 | -3.165 | 1.094 | **-5.469** | **-1.194** | 1 | 5421.948 | 6007.998 |

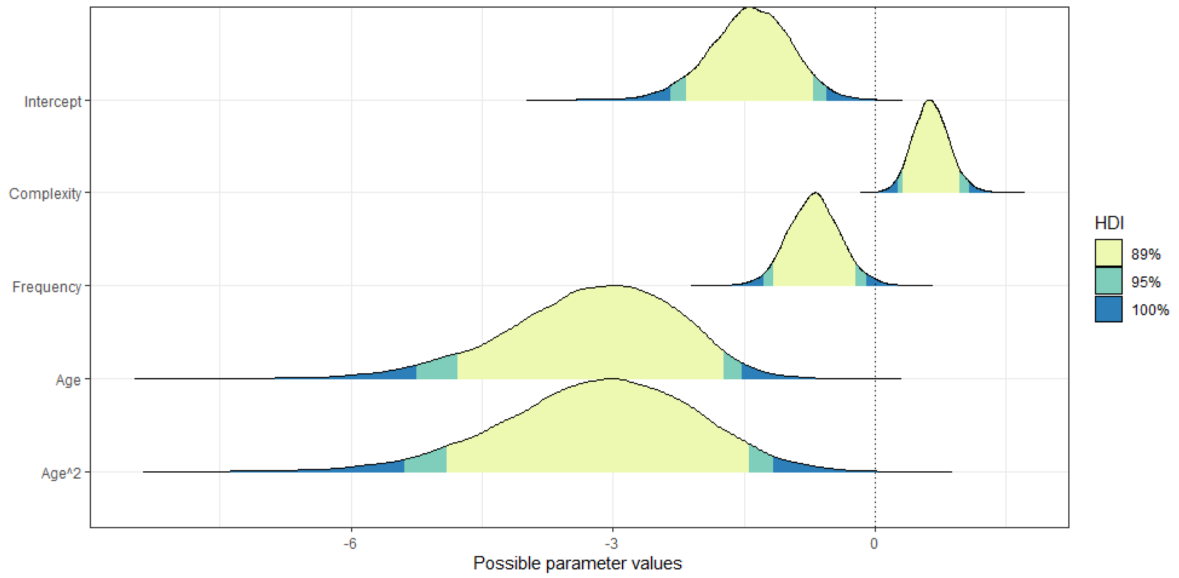


Figure S2: HDI of Model A, related to Figure 2. HDI are shown for each model estimate with credibility intervals of 89% marked in yellow, of 95% marked in green, and of 100% marked in blue. All model predictors were z-transformed.

Table S2: Supplementary Model A.1 output, related to STAR Methods Model A. Model estimates and respective standard error, credibility intervals (Q2.5-Q97.5), Rhats, Bulk ESS and Tail ESS. All predictors (“complexity”, “frequency”, “age”, “$age^2$”) were z-transformed. Credibility intervals not comprising zero are marked in bold.

| | Estimate | Est.Error | Q2.5 | Q97.5 | Rhat | Bulk_ESS | Tail_ESS |
|---|---|---|---|---|---|---|---|
| Intercept | -1.448 | 0.466 | **-2.426** | **-0.599** | 1.001 | 10093.531 | 9659.126 |
| comp_z | 0.737 | 0.33 | **0.113** | **1.415** | 1 | 8415.279 | 8225.436 |
| freq_z | -0.685 | 0.31 | **-1.291** | **-0.057** | 1 | 9775.662 | 8926.204 |
| age_z | -3.289 | 1.004 | **-5.481** | **-1.598** | 1 | 9490.216 | 7153.911 |
| age_z2 | -3.213 | 1.13 | **-5.644** | **-1.208** | 1 | 9020.363 | 6985.53 |
| comp_z: freq_z | 0.121 | 0.344 | -0.549 | 0.821 | 1 | 8971.779 | 9114.571 |

Table S3: Supplementary Model A.2 output, related to STAR Methods Model A. Model estimates and respective standard error, credibility intervals (Q2.5-Q97.5), Rhats, Bulk ESS and Tail ESS. All predictors (“complexity”, “frequency”, “age”, “$age^2$”) were z-transformed. Credibility intervals not comprising zero are marked in bold.

| | Estimate | Est.Error | Q2.5 | Q97.5 | Rhat | Bulk_ESS | Tail_ESS |
|---|---|---|---|---|---|---|---|
| Intercept | -1.717 | 0.506 | **-2.775** | **-0.804** | 1 | 6444.141 | 8128.623 |
| comp_z | 0.828 | 0.339 | **0.18** | **1.51** | 1 | 4813.061 | 7484.446 |
| age_z | -3.95 | 1.186 | **-6.697** | **-2.038** | 1.001 | 4030.568 | 4661.856 |
| age_z2 | -3.563 | 1.203 | **-6.245** | **-1.543** | 1.001 | 4022.643 | 5005.549 |
| freq_z | -0.818 | 0.444 | -1.725 | 0.028 | 1 | 5217.621 | 6586.852 |
| comp_z: age_z | 0.065 | 0.927 | -1.717 | 1.939 | 1.001 | 4142.497 | 4919.346 |
| comp_z: age_z2 | -0.429 | 0.937 | -2.242 | 1.435 | 1.001 | 4510.801 | 5338.179 |

| | | | | | | | |
|---|---|---|---|---|---|---|---|
| age_z: freq_z | 0.085 | 1.027 | -2.149 | 1.963 | 1 | 4102.082 | 4153.779 |
| age_z2: freq_z | 0.419 | 1.015 | -1.718 | 2.316 | 1 | 4208.113 | 4682.544 |

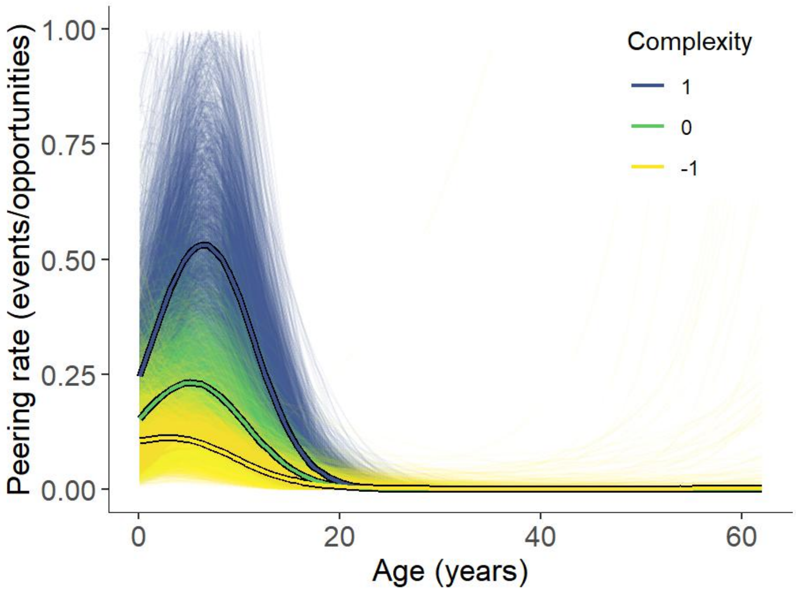


Figure S3: Peering frequencies at foot item complexity, related to Figure 2. Relative peering frequencies at different food item complexities (z-transformed mean complexity +/- SD) over age, using model estimates from Supplementary Model A.2.

<u>**Model B:** Effect of target age on peering frequencies across development</u>

Table S4: Model B output, related to Figure 3. Model estimates and respective standard error, credibility intervals (Q2.5-Q97.5), Rhats, Bulk ESS and Tail ESS; continuous predictors (age, $age^2$) were z-transformed; categorical predictor age relation was dummy coded (reference category age mate). Credibility intervals not comprising zero are marked in bold.

| | Estimate | Est.Error | Q2.5 | Q97.5 | Rhat | Bulk_ESS | Tail_ESS |
|---|---|---|---|---|---|---|---|
| Intercept | -12.017 | 9.04 | **-37.469** | **-1.844** | 1.003 | 969.523 | 609.514 |
| age_z | -44.551 | 37.322 | **-149.34** | **-4.592** | 1.003 | 958.341 | 567.904 |
| age_z2 | -44.717 | 37.615 | **-150.965** | **-7.034** | 1.003 | 970.283 | 564.147 |
| AgeRelation Older | 11.22 | 9.041 | **1.069** | **36.675** | 1.003 | 971.636 | 606.436 |
| AgeRelation Younger | 8.585 | 9.048 | -1.655 | 34.127 | 1.003 | 968.333 | 605.911 |
| age_z: AgeRelation Older | 42.782 | 37.318 | **2.789** | **147.704** | 1.003 | 959.546 | 567.54 |
| age_z: AgeRelation Younger | 43.773 | 37.336 | **3.809** | **149.235** | 1.003 | 959.51 | 565.39 |
| age_z2: AgeRelation Older | 43.377 | 37.612 | **5.673** | **149.638** | 1.003 | 970.709 | 564.636 |
| age_z2: AgeRelation Younger | 44.785 | 37.616 | **7.028** | **151.237** | 1.003 | 970.092 | 560.705 |

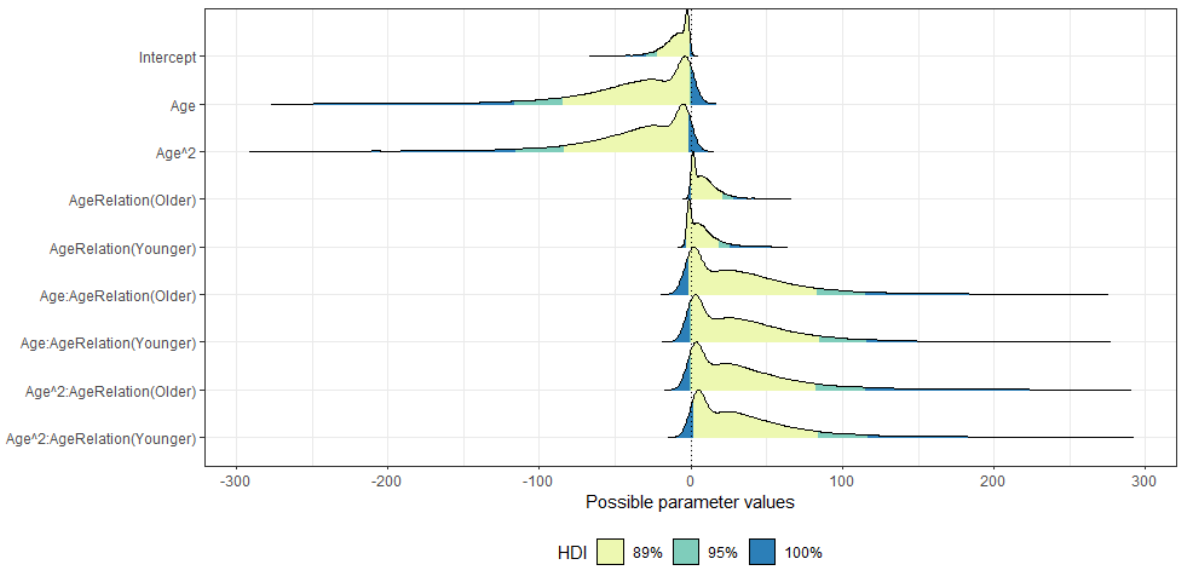


Figure S4: HDI Model B, related to Figure 3. HDI are shown for each model estimate with credibility intervals of 89% marked in yellow, of 95% marked in green, and of 100% marked in blue; continuous predictors (age, age$^2$) were z-transformed; categorical predictor age relation was dummy coded (reference category age mate).

**<u>Model C1:</u>** <u>Effect of kin relation on overall peering frequencies</u>

Table S5: Model C1 output, related to Figure 4. Model estimates and respective standard error, credibility intervals (Q2.5-Q97.5), Rhats, Bulk ESS and Tail ESS; continuous predictors (age, age$^2$) were z-transformed; categorical predictor kin relation was dummy coded (reference category unrelated). Credibility intervals not comprising zero are marked in bold.

| | Estimate | Est.Error | Q2.5 | Q97.5 | Rhat | Bulk_ESS | Tail_ESS |
|---|---|---|---|---|---|---|---|
| Intercept | -1.765 | 0.338 | **-2.449** | **-1.114** | 1 | 5146.806 | 7071.504 |
| age_z | -0.056 | 0.395 | -0.878 | 0.681 | 1 | 4647.958 | 5209.917 |
| age_z2 | -0.357 | 0.242 | -0.84 | 0.125 | 1 | 4929.654 | 6411.342 |
| target Mother | -1.147 | 0.392 | **-1.935** | **-0.39** | 1 | 7425.197 | 8171.355 |
| target Kin | -2.759 | 0.689 | **-4.264** | **-1.55** | 1 | 5121.764 | 6057.572 |
| age_z: target Mother | -3.452 | 0.976 | **-5.717** | **-1.93** | 1 | 5108.345 | 4644.693 |
| age_z: target Kin | -6.372 | 2.769 | **-12.333** | **-1.759** | 1 | 3526.883 | 4382.222 |
| age_z2: target Mother | -0.457 | 0.998 | -2.808 | 1.02 | 1.001 | 5488.403 | 4459.734 |
| age_z2: target Kin | -5.232 | 2.787 | **-11.231** | **-0.493** | 1 | 3857.775 | 4906.911 |

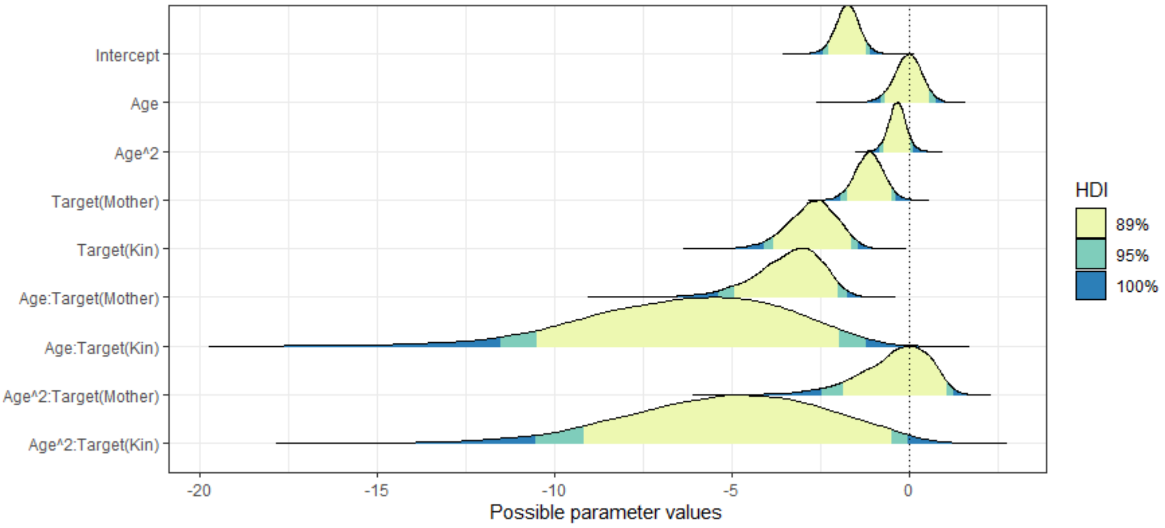


Figure S5: HDI of Model C1, related to Figure 4. HDI are shown for each model estimate with credibility intervals of 89% marked in yellow, of 95% marked in green, and of 100% marked in blue; continuous predictors (age, $age^2$) were z-transformed; categorical predictor kin relation was dummy coded (reference category unrelated).

**<u>Model C2:</u>** <u>Effect of kin relation on peering opportunity use</u>

Table S6: Model C2 output, related to Figure 4. Model estimates and respective standard error, credibility intervals (Q2.5-Q97.5), Rhats, Bulk ESS and Tail ESS; continuous predictors (age, $age^2$) were z-transformed; categorical predictor kin relation was dummy coded (reference category unrelated). Credibility intervals not comprising zero are marked in bold.

| | Estimate | Est.Error | Q2.5 | Q97.5 | Rhat | Bulk_ESS | Tail_ESS |
|---|---|---|---|---|---|---|---|
| Intercept | -0.536 | 0.295 | -1.11 | 0.046 | 1.001 | 6673.892 | 7721.3 |
| age_z | 0.302 | 0.364 | -0.422 | 1.016 | 1.001 | 7420.319 | 8226.291 |
| age_z2 | -0.418 | 0.18 | **-0.79** | **-0.076** | 1 | 7421.844 | 8173.335 |
| target Mother | -0.997 | 0.319 | **-1.638** | **-0.39** | 1 | 12234.16 | 9283.263 |
| target Kin | -2.365 | 0.515 | **-3.428** | **-1.394** | 1.001 | 13320.424 | 8299.313 |
| age_z: target Mother | -2.112 | 0.864 | **-4.024** | **-0.713** | 1.001 | 7099.669 | 6020.777 |
| age_z: target Kin | -5.383 | 2.024 | **-9.822** | **-2.059** | 1 | 5758.19 | 5715.2 |
| age_z2: target Mother | -0.748 | 0.953 | -2.925 | 0.719 | 1 | 7830.768 | 5951.09 |
| age_z2: target Kin | -4.058 | 2.521 | -9.476 | 0.13 | 1 | 6239.661 | 5763.652 |

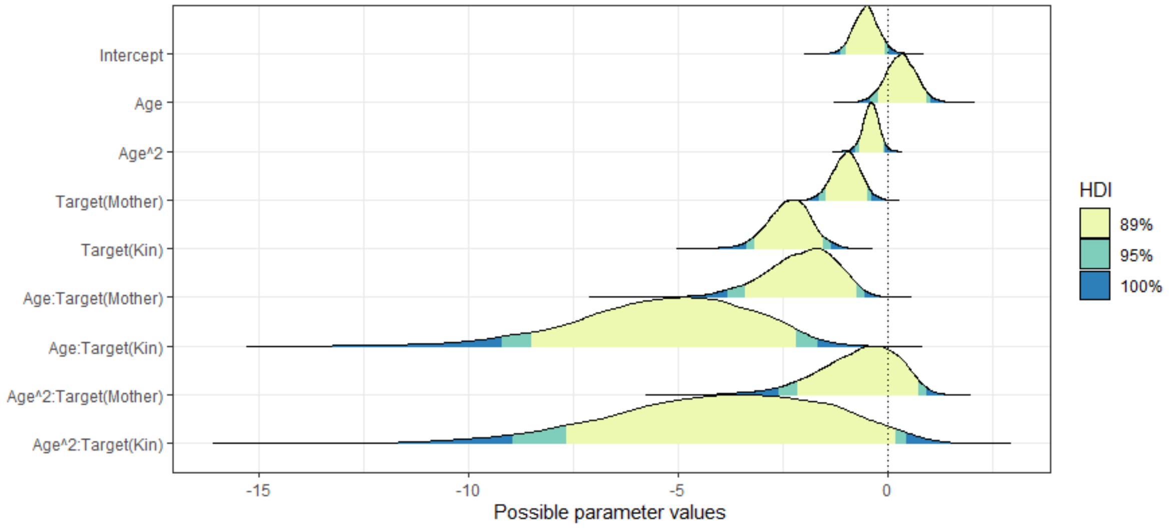


Figure S6: HDI of Model C2. HDI are shown for each model estimate with credibility intervals of 89% marked in yellow, of 95% marked in green, and of 100% marked in blue; continuous predictors (age, age$^2$) were z-transformed; categorical predictor kin relation was dummy coded (reference category unrelated).

<u>**Model D:** Effect of conspecific dominance rank on peering frequencies</u>

Table S7: Model D output. Model estimates and respective standard error, credibility intervals (Q2.5-Q97.5), Rhats, Bulk ESS and Tail ESS; continuous predictors (age, age$^2$) were z-transformed; categorical predictors rank and sex were dummy coded (reference categories high, female). Credibility intervals not comprising zero are marked in bold.

| | Estimate | Est.Error | Q2.5 | Q97.5 | Rhat | Bulk_ESS | Tail_ESS |
|---|---|---|---|---|---|---|---|
| Intercept | -1.786 | 0.376 | **-2.55** | **-1.054** | 1.001 | 9825.808 | 9614.958 |
| age_z | -2.391 | 0.517 | **-3.497** | **-1.465** | 1 | 10138.487 | 7616.082 |
| age_z2 | -1.699 | 0.596 | **-2.949** | **-0.6** | 1 | 9145.943 | 7078.134 |
| sex male | 0.446 | 0.384 | -0.314 | 1.202 | 1 | 11803.825 | 9982.896 |
| rank low | -0.049 | 0.354 | -0.763 | 0.634 | 1 | 12902.573 | 9698.646 |
| rank medium | -0.204 | 0.3 | -0.789 | 0.382 | 1 | 11712.323 | 9321.849 |
| sex male: rank low | -0.741 | 0.731 | -2.178 | 0.655 | 1 | 13384.67 | 10177.739 |
| sex male: rank medium | 0.611 | 0.519 | -0.394 | 1.636 | 1 | 10702.362 | 9079.719 |

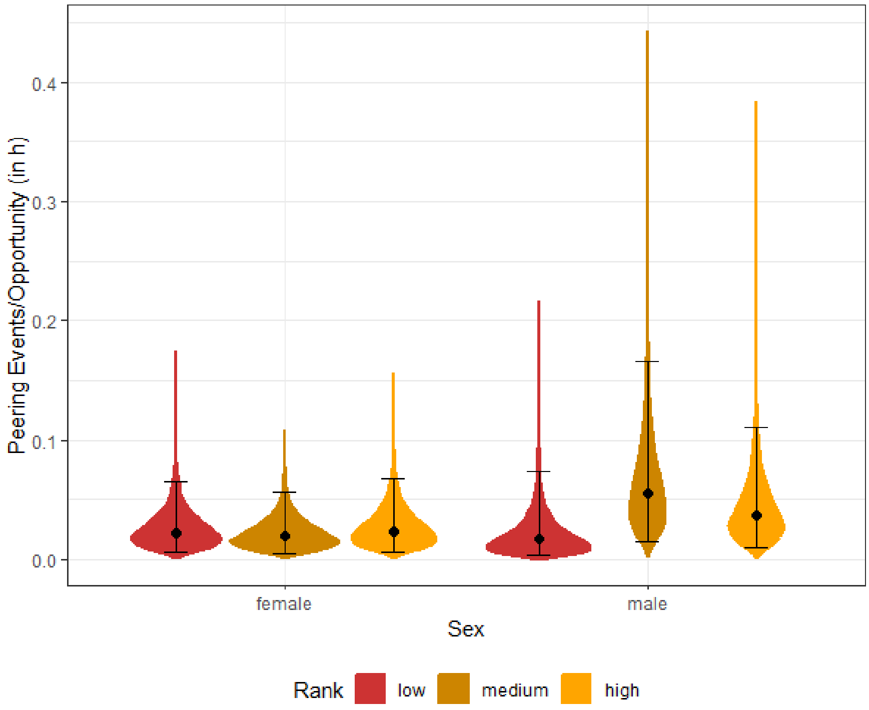


Figure S7: Model Predictions of Model D. Relative peering frequencies directed at low (red), medium (brown), and high (orange) ranking females (left) and males (right).

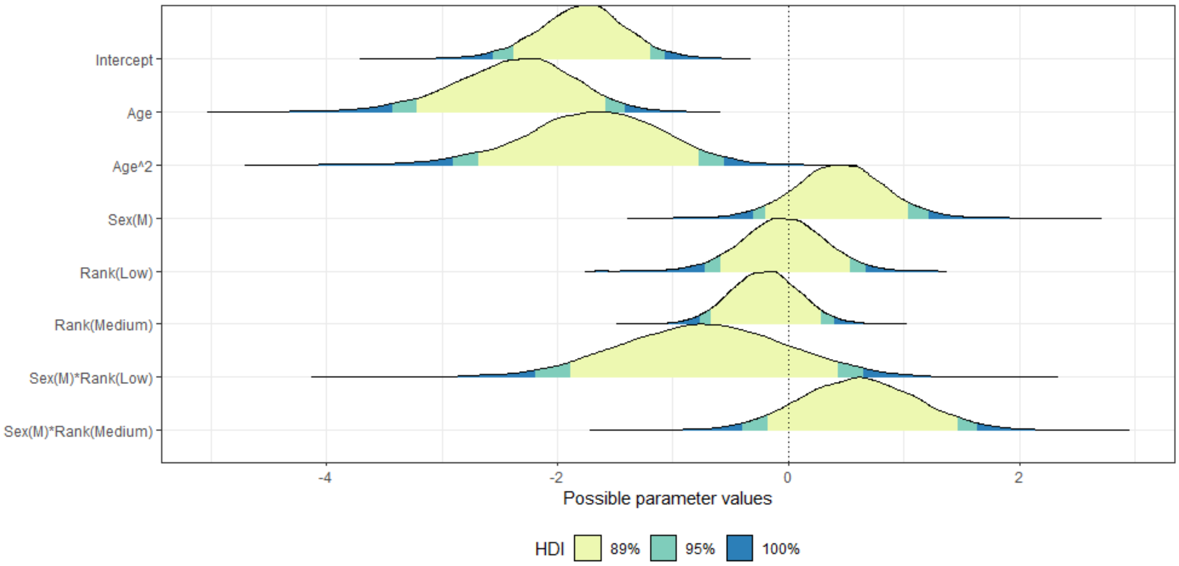


Figure S8: HDI of Model D. HDI are shown for each model estimate with credibility intervals of 89% marked in yellow, of 95% marked in green, and of 100% marked in blue; continuous predictors (age, age$^2$) were z-transformed; categorical predictors rank and sex were dummy coded (reference categories high, female).

**Model E:** Increased daily grooming rates on peering days

Table S8: Model E output. Model estimates and respective standard error, credibility intervals (Q2.5-Q97.5), Rhats, Bulk ESS and Tail ESS; categorical predictor peering was dummy coded (reference category no). Credibility intervals not comprising zero are marked in bold.

| | Estimate | Est.Error | Q2.5 | Q97.5 | Rhat | Bulk_ESS | Tail_ESS |
|---|---|---|---|---|---|---|---|
| Intercept | -2.167 | 0.104 | **-2.37** | **-1.965** | 1.002 | 1977.584 | 3640.635 |
| zi_Intercept | -1.427 | 0.229 | **-1.942** | **-1.034** | 1 | 8900.214 | 10449.209 |
| peerBin2yes | 0 | 0.086 | -0.169 | 0.17 | 1 | 14663.57 | 9898.222 |
| zi_peerBin2yes | -14 | 10.375 | **-41.845** | **-2.933** | 1 | 6132.995 | 4738.023 |

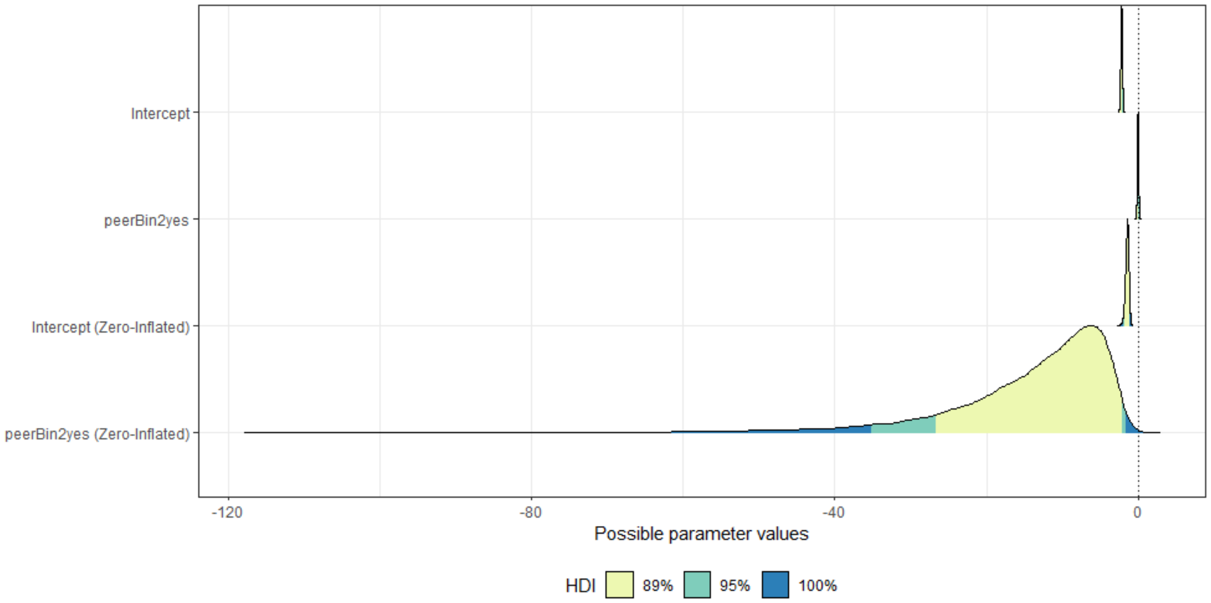


Figure S9: HDI of Model E. HDI are shown for each model estimate with credibility intervals of 89% marked in yellow, of 95% marked in green, and of 100% marked in blue; categorical predictor peering was dummy coded (reference category no).

**Peering across different contexts**

Table S9: Observed behavioral repertoire. List of single behaviors of observed behavioral repertoire by category (A=Feeding in green, B=Grooming in light purple, C=Social in dark purple, D=Exploration in light green, E=Object use in olive, F=Other in dark green) with definition, previous recognition of cultural behavior, and observed occurrences during scan sampling (Focal or Close Party Member) and all occurrence sampling of Peering (only focal individual).

| | Name of behavior | Definition | Previous Cultural Recognition | Occurrences | | |
|---|---|---|---|---|---|---|
| | | | | Focal | Close Party Member | Peering |
| **A.** | **Feeding** | **Focal individual engages in proficient feeding behavior, including possibly detachment of food item, possibly processing of food item, and feeding (i.e. chewing).** | **-** | **6185** | **6258** | **178** |
| **1.** | ***Bark*** | Focal individual feeds on bark of a plant. | | | | |
| | AB | *Alstonia boonei.* | - | 1 | 6 | 1 |
| | CGP | *Celtis durandii/gomphophylla* | - | 1 | - | 0 |
| | CHA | *Chaetachme aristata.* | - | - | 1 | 0 |
| | CP | *Cleistopholis patens.* | - | 8 | 27 | 2 |
| | CYA | *Cynometra alexandrii* | - | - | 3 | 0 |
| | CZE | *Celtis zenkeri* | - | 1 | - | 0 |
| | PSG | *Psidium guajava* | - | 1 | - | 0 |
| | THV | General terrestrial herbaceous vegetation, regardless of species. | - | - | - | 1 |
| | TRR | *Trichilia rubescens* | - | 1 | - | 0 |
| | Unidentified species 1 | Unidentified swamp tree species. | - | - | - | 1 |
| **2.** | ***Flowers*** | Focal individual feeds on flowers of a plant. | | | | |
| | BPY | *Broussonetia papyrifera* | - | 262 | 198 | 0 |
| | CLI | General climber, regardless of species. | - | 9 | 22 | 0 |
| | CMI | *Celtis mildbraedii* | - | 72 | 59 | 0 |
| | LP | *Lasiodiscus mildbraedii* | - | 1 | 10 | 1 |
| | URC | *Urera camaroonensis/trinervis* | - | 33 | 21 | 0 |
| **3.** | ***Fruits*** | Focal individual feeds on fruits of a plant. | | | | |

| | | | | | | |
|---|---|---|---|---|---|---|
| | AFM | *Aframomum* sp. | - | 6 | 6 | 1 |
| | ALP | *Alaphia* sp. | - | 80 | 35 | 6 |
| | ANM | *Antrocarium micrantha* | - | 61 | 19 | 0 |
| | ANT | *Antiaris toxicaria* | - | 72 | 160 | 0 |
| | BEO | *Bequaerhodendron oblanceolatum* | - | 8 | 16 | 0 |
| | BPY | *Broussonetia papyrifera* | - | 147 | 157 | 1 |
| | CGO | *Chrysophyllum gorungosanum* | - | 72 | 36 | 0 |
| | CGP | *Celtis durandii/gomphophylla* | - | 59 | 5 | 0 |
| | CLI | General climber, regardless of species. | - | 4 | 9 | 3 |
| | CLS | *Caloncoba schweinfurthii* | - | 3 | - | 0 |
| | COG | *Cola gigantea* | - | 6 | - | 0 |
| | COM | *Cordia millenii* | - | 189 | 118 | 3 |
| | CRA | *Crossonephelis africanus* | - | - | 28 | 1 |
| | CYA | *Cynometra alexandrii* | - | 120 | 178 | 2 |
| | DD | *Desplatsia dewevrei* | - | 72 | 42 | 10 |
| | EKC | *Ekebergia capensis* | - | 3 | - | 0 |
| | ES | *Erythrophleum suaveolens* | - | 13 | - | 0 |
| | FB | *Ficus barteri* | - | 108 | 46 | 0 |
| | FE | *Ficus exasperata* | - | 330 | 234 | 9 |
| | FIC | *Ficus* spp. | - | 9 | 1 | 1 |
| | FM | *Ficus mucuso* | - | 1285 | 1268 | 18 |
| | FN | *Ficus natalensis* | - | 140 | 169 | 0 |
| | FO | *Ficus ottomaefolia* | - | 5 | - | 0 |
| | FPO | *Ficus polita* | - | 8 | 6 | 0 |
| | FSA | *Ficus sansibarica (brachylepsis)* | - | 2 | - | 0 |
| | FSS | *Ficus saussureana (dawei/lutea)* | - | 5 | 10 | 0 |
| | FSU | *Ficus sur (capensis/vogelana)* | - | 644 | 1475 | 7 |
| | FVL | *Ficus vallis-choudae* | - | 2 | - | 0 |
| | FVR | *Ficus variifolia* | - | 135 | 81 | 1 |
| | GAL | *Chysophyllum albidum* | - | 9 | 37 | 2 |
| | GPR | *Chrysophyllum perpulchrum* | - | 5 | 5 | 0 |
| | KLG | *Klainedoxa gabonensis* | - | 16 | 26 | 2 |

| | | | | | | |
|---|---|---|---|---|---|---|
| | LCA | *Lantana camara* | - | 4 | - | 0 |
| | MA | *Mammea africana* | - | 49 | 59 | 11 |
| | MEX | *Milicia (Chlorophora) excelsa* | - | 109 | 27 | 0 |
| | MIE | *Mildbraediodendron excelsum* | - | 69 | 43 | 15 |
| | MMA | *Mango mangifera* | - | 85 | 220 | 0 |
| | MOL | *Morus lactea/mesozygia* | - | 9 | 16 | 0 |
| | MYH | *Myrianthus holstii* | - | 13 | 42 | 11 |
| | PF | *Parkia filicoidea* | - | 2 | 5 | 0 |
| | PSG | *Psidium guajava* | - | 32 | 1 | 0 |
| | PSM | *Pseudospondias microcarpa* | - | 11 | 6 | 5 |
| | RF | *Raphia farinifera* | - | 5 | 8 | 0 |
| | SAB | *Saba florida* | - | - | 1 | 2 |
| | STD | *Sterculia dawei* | - | 7 | 1 | 0 |
| | TRA | *Treculia africana* | - | 39 | 136 | 17 |
| | Unidentified species 2 | Unidentified climber species with small orange fruits. | - | 2 | - | 0 |
| **4.** | ***Insects*** | Focal individual feeds on insects or insect products. | | | | |
| | Honey | Focal individual feeds on honey, regardless of insect species. | - | 17 | 12 | 0 |
| | Termite soil | Focal individual feeds on termite soil. | - | 13 | 9 | 8 |
| **5.** | ***Leaves*** | Focal individual feeds on leaves of a plant. | | | | |
| | AM | *Argomuellera macrophylla* | - | 1 | - | 0 |
| | ANT | *Antiaris toxicaria* | - | 9 | 7 | 0 |
| | BPY | *Broussonetia papyrifera* | - | 420 | 222 | 5 |
| | CGP | *Celtis durandii/gomphophylla* | - | 10 | - | 0 |
| | CHA | *Chaetachme aristata* | - | 1 | 2 | 0 |
| | CLI | *Climber general* | - | 29 | 17 | 1 |
| | CLS | *Caloncoba schweinfurthii* | - | 1 | - | 0 |
| | CMI | *Celtis mildbraedii* | - | 366 | 209 | 2 |
| | CPH | *Celtis wightii/philippensis* | - | 193 | 130 | 1 |
| | CYA | *Cynometra alexandrii* | - | 5 | 8 | 0 |
| | CZE | *Celtis zenkeri* | - | 15 | 15 | 1 |
| | DD | *Desplatsia dewevrei* | - | 5 | - | 0 |

| | | | | | | |
|---|---|---|---|---|---|---|
| | EPI | Epiphyte (tree cabbage) | - | - | - | 1 |
| | FE | *Ficus exasperata* | - | 203 | 128 | 2 |
| | FVR | *Ficus variifolia* | - | 84 | 54 | 1 |
| | LM | *Lasiodiscus mildbraedii* | - | 3 | 22 | 0 |
| | LT | *Lovoa trichilioides* | - | 3 | - | 0 |
| | MOL | *Morus lactea/mesozygia* | - | 1 | 1 | 0 |
| | STD | *Sterculia dawei* | - | - | 1 | 0 |
| | THV | Terrestrial herbaceous vegetation | - | 3 | 5 | 0 |
| | TRD | *Trichilia dregeana* | - | 6 | 4 | 1 |
| | URC | *Urera camaroonensis/trinervis* | - | 3 | 2 | 0 |
| **6.** | ***Meat*** | Focal individual feeds on meat. | | | | |
| | BD | Blue duiker (*Philantomba monticola*) | - | 10 | - | 0 |
| | BM | Blue monkey (*Cercopithecus mitis*) | - | 4 | - | 0 |
| | BWC | Black and white colobus (*Colobus guereza)* | - | 85 | 120 | 13 |
| **7.** | ***Pith*** | Focal individual feeds on pith, regardless of plant species. | - | 58 | 49 | 3 |
| **8.** | ***Resin*** | Focal individual feeds on resin of a plant. | | | | |
| | KA | *Khaya anthoteca* | - | 136 | 69 | 2 |
| | RF | *Raphia farinifera* | - | 6 | 21 | 0 |
| **9.** | ***Roots*** | Focal individual feeds on roots, regardless of plant species. | - | 7 | 14 | 0 |
| **10.** | **Soil** | Focal individual feeds on soil. | - | 5 | 8 | 0 |
| **11.** | ***Water*** | Focal individual drinks water without using a tool. | - | 3 | - | 1 |
| **12.** | ***Wood*** | Focal individual feeds on wood (fresh or dead), regardless of plant species. | - | 36 | 43 | 2 |
| **B.** | **Grooming** | **Focal individual engages in grooming, i.e. removal of dirt, defoliated skin, ectoparasites from own fur or fur of conspecifics.** | | **2392** | **3974** | **92** |
| **1.** | ***Self*** | Focal individual grooms own body part. | - | 867 | 682 | 32 |
| **2.** | ***Self wound*** | Focal individual grooms wound, i.e. touches wound, picks at wounds, licks wound. Here, exclusively own wound. | - | 1 | 3 | 6 |

| | | | | | | |
|---|---|---|---|---|---|---|
| **3.** | ***Social*** | Focal individual grooms or is being groomed by a conspecific. | - | 1523 | 3289 | 54 |
| **4.** | ***Social hand-clasp*** | Focal individual grooms or is being groomed by conspecific while they clasp hands or wrists above their heads. (1) | - | 1 | - | 0 |
| **C.** | **Social** | **Focal individual engages in social activity with other conspecific(s), excluding social grooming (see above), or displays behavior towards conspecific(s).** | | **1707** | **2298** | **30** |
| **1.** | ***Cling*** | Individual, here immature, is holding on to conspecific, usually the mother. | | | | |
| | ***Resting on back*** | Immature sitting on and/or holding on to the mothers back, commonly while the mother is travelling. | - | 237 | 366 | 0 |
| | ***Resting in cling*** | Immature is clinging to the belly or side of the mother, being carried during travels or other activities. (2) | - | 818 | 776 | 3 |
| | ***Resting while holding onto conspecific*** | Immature rests while holding onto conspecific, usually the mother. | - | 10 | 11 | 0 |
| **2.** | ***Social agonistic*** | Focal individual engages in aggressive behavior with other conspecific(s). | | | | |
| 2.1. | Agonistic | Focal individual engages in a general social agonistic behavior. Not covered by chase, displacement, flee, severe (see below). | - | - | 3 | 1 |
| 2.2. | Chase | Focal individual chases other conspecific over more than 10 m or into another tree. (2) | - | 2 | 4 | 0 |
| 2.3. | Displacement | Focal individual leaves because another conspecific approaches. | - | 1 | 1 | 0 |
| 2.4. | Flee | Focal individual retreats fast from another conspecific. (2) | - | 3 | 3 | 0 |
| 2.5 | Severe | Focal individual engages in aggressive behavior involving body contact with conspecific(s). | - | 3 | 1 | 0 |
| **3.** | ***Approach*** | Focal individual cautiously approaches another conspecific, observing them attentively. | - | 5 | 6 | 1 |

| | | | | | | |
|---|---|---|---|---|---|---|
| **4.** | ***Beg*** | Focal individual begs by reaching or moving mouth toward the hand or mouth of a conspecific. (2) | - | 3 | 10 | 4 |
| **5.** | ***Collect*** | "Mother actively collects the infant into body contact." (2) | - | 1 | 11 | 0 |
| **6.** | ***Copulation*** | Focal individual copulates with conspecific. | - | 2 | 2 | 4 |
| **7.** | ***Display*** | Focal individual engages in display behavior toward a conspecific. (2) | - | 6 | 5 | 1 |
| **8.** | ***Embrace*** | Focal individual touches the shoulder/back of another conspecific. (2) | - | 2 | 3 | 0 |
| **9.** | ***Sex*** | Focal individual engages in sexual interactions with other conspecific(s). | | | | |
| 9.1. | Inspect | "Focal individual sniffs or touches with mouth or fingers the vagina/penis of other conspecific." (2) | - | 8 | 5 | 2 |
| 9.2. | Present | Focal individual approaches another conspecific, directing genitals toward them. (3) | - | 1 | 4 | 0 |
| 9.3. | Stimulate | Focal individual self-stimulates genitals with hands or tools. (2,3) | - | 1 | - | 0 |
| **10.** | ***Listen*** | Focal individual listens actively to vocalizations from conspecific(s), whether from the same or different party. | - | 15 | 6 | 0 |
| **11.** | ***Play*** | Focal individual engages in social play with other conspecific(s). (2) | - | 435 | 897 | 7 |
| **12.** | ***Peer*** | Focal individual peers at other conspecific(s). | - | 90 | 130 | 1 |
| **13.** | ***Touch*** | Focal individual touches another conspecific. | - | 1 | 6 | 1 |
| **14.** | ***Social vocalization*** | Focal produces directed or undirected call. | | | | |
| 14.1. | Greet | Focal individual emits a pant grunt (a series of joined grunts) as a submissive signal to a higher-ranking conspecific. (4) | - | 6 | 8 | 1 |
| 14.2. | Vocalization | Focal individual emits a general vocalization not included in the categories listed. | - | - | 4 | 2 |

| | | | | | | |
|---|---|---|---|---|---|---|
| 14.3. | Alarm call | Focal individual emits this type of vocalization to signal the presence of danger to other conspecific(s). (5) | - | 6 | 6 | 0 |
| 14.4. | Hoos | Focal individual emits this type of vocalization "during feeding, as a sign of mild fear or surprise, or when travelling near nesting time." (4) | - | - | 3 | 0 |
| 14.5. | Hoot | Focal individual emits this type of vocalization during an inter-group encounter or as a "long distance cohesive calling between temporary groups." (6) | - | - | 5 | 0 |
| 14.6. | Panthoot | Focal individual emits this type of vocalization "during feeding, travelling, meeting chimpanzees from other parties or other communities." (4) | - | 3 | 1 | 1 |
| 14.7. | Scream | Focal individual emits this type of vocalization "when receiving aggression, as a sign of fear, as part of pant hoot or during a copulation." (4) | - | - | 1 | 0 |
| 14.8. | Tantrum | Immature individual engages in repetitive movements, often accompanied by screaming or crying. (2) | - | - | - | 1 |
| 14.9. | Whimper | "Infants and juveniles emit this type of vocalization during weaning or separation from the mother or by young adult males when separated from other males." (4) | - | - | 1 | 0 |
| **15.** | ***Social waiting*** | Focal individual waits for a conspecific, usually in the context of a mother and infant pair. (2) | - | 13 | 3 | 0 |
| **16.** | ***Social watching*** | Focal individual looks attentively at a conspecific more than 5 meters away. | - | 33 | 14 | 0 |
| **1z.** | ***Social watching of vocalizing party*** | Focal individual looks attentively at other conspecifics vocalizing from over 5 meters away. | - | 2 | 2 | 0 |
| **D.** | **Exploratory behaviors** | **Focal individual engages in attentive manipulation of, inspection of and/or play with an object/the immediate environment.** | | **677** | **554** | **17** |
| **1.** | ***Auto play*** | Focal individual engages in solitary playful behavior. Behavior is often repetitive and may include practice of skills. (2) | | | | |

| | | | | | | |
|---|---|---|---|---|---|---|
| 1.1. | Display | Focal individual engages in "'Playful display' with branches, usually by immatures." (2) | - | 1 | - | 0 |
| 1.2. | Movement | Focal individual engages in "repetitive movement such as twirling, swinging, etc., also repetitive swinging of just one arm or leg." (2) | - | 1<br>38 | 190 | 0 |
| 1.3. | Nesting | "Infant makes or tries to make nests." (2) | - | 9 | 2 | 0 |
| 1.4. | Object | Focal individual engages in "manipulation of objects with no apparent immediate goal, including repetitive movements with objects." (2) | - | 75 | 40 | 1 |
| **2.** | ***Exploration (General)*** | Focal individual engages in attentive manipulation and/or inspection of an object/the immediate environment. | - | 101 | 60 | 8 |
| **3.** | ***Food search*** | Focal individual engages in exploration that results in feeding, e.g. searching fruits on ground and feeding on fruits after, digging for roots and feeding on roots after. | - | 19 | 14 | 6 |
| **4.** | ***Try feeding*** | Focal individual "attempts to feed on a food item or other object whereby the item is taken into the mouth but not properly processed and not ingested." (2) | - | 334 | 248 | 2 |
| **E.** | **Object Use** | **Focal engages in tool use, object use, or construction behavior. (7)** | | **86** | **42** | **20** |
| **1.** | ***Nesting*** | Focal individual uses multiple attached branches and twigs and by intertwining creates a (semi)permanent nest structure. (8) | - | 23 | 15 | 3 |
| **2.** | ***Seat-vegetation*** | Focal individual sits on leaves placed on the ground, leaves are either detached or gathered by the individual. (8) | Yes | 1 | 1 | 0 |
| **3.** | ***Leaf clip*** | Focal individual noisily rips a leaf using mouth or fingers to gain attention for various functions. (8) | Yes | 3 | - | 0 |
| **4.** | ***Ectoparasite inspection on leaf*** | Focal individual inspects and/or squashes ectoparasites on leaves. Previously defined as leaf-squash and leaf-inspect. (8) | Yes | 17 | 11 | 12 |

| | | | | | | |
|---|---|---|---|---|---|---|
| **5.** | ***Leaf sponge*** | Focal individual squishes/chews on bundle of leaves (making a leaf sponge), dips leaf sponge in water and drinks from it. (8) | Candidate | 36 | 13 | 4 |
| **6.** | ***Leaf napkin*** | Focal individual uses leaves to clean body parts. (9) | Yes | 2 | - | 0 |
| **7.** | ***Scratch*** | Focal individual manipulates an object (e.g., branch) for self-scratching. | - | 4 | 1 | 0 |
| **8.** | ***Leaf dab wound*** | Focal individual touches/presses leaf on wound, possibly after wetting leaf with saliva, and possibly inspects leaf after. (8) | Yes | - | 1 | 1 |
| **F.** | **Other** | **Focal individual engages in behavior not fitting the contexts listed above.** | | **6536** | **8414** | **7** |
| **1.** | ***Drinking*** | Infant drinks milk from the mother | - | 107 | 35 | 0 |
| **2.** | ***Hunting*** | Focal individual engages in hunting behavior. | - | 18 | 4 | 0 |
| **3.** | ***Moving*** | Focal individual moves within or between trees. | - | 771 | 655 | 2 |
| **4.** | ***Moving go to*** | Focal individual, commonly an infant, moves through trees toward another conspecific, usually the mother. | - | 47 | 27 | 0 |
| **5.** | ***Resting*** | Focal individual sits or lies down, not visibly engaging in any active behavior. | - | 3425 | 5237 | 4 |
| 5.1. | Defecate | Focal individual defecates while sitting or lying down, not visibly engaging in any active behavior. | - | 4 | - | 0 |
| 5.2. | Urinate | Focal individual urinates while sitting or lying down, not visibly engaging in any active behavior. | - | 14 | 8 | 0 |
| 5.3. | Nest | Focal individual sits or lies down in a nest, not visibly engaging in any active behavior. | - | 870 | 932 | 0 |
| 5.4. | Nest urinate | Focal individual urinates while sitting or lying down in a nest, not visibly engaging in any active behavior. | - | 1 | 1 | 0 |
| **6.** | ***Travelling*** | Focal individual walks on ground. | - | 1267 | 1493 | 1 |
| **7.** | ***Travelling go to*** | Focal individual, commonly an infant, walks on the ground toward another conspecific, usually the mother. | - | 12 | 22 | 0 |
